# Endocytosis is regulated through the pH-dependent phosphorylation of Rab GTPases by Parkinson’s kinase LRRK2

**DOI:** 10.1101/2023.02.15.528749

**Authors:** Michelle E. Maxson, Kassidy K. Huynh, Sergio Grinstein

## Abstract

While it has been known for decades that luminal acidification is required for normal traffic along the endocytic pathway, the precise underlying mechanism(s) remain unknown. We found that dissipation of the endomembrane pH gradient resulted in acute formation of large Rab5- or Rab7-positive vacuoles. Vacuole formation was associated with and required hyperactivation of the Rabs, which was attributable to impaired GTPase activity, despite normal recruitment of cognate GAPs. Surprisingly, LRRK2 –a kinase linked to Parkinson’s disease–was recruited to endomembranes and markedly activated upon dissipation of luminal acidification. LRRK2 phosphorylated Rab GTPases, rendering them insensitive to deactivation. Importantly, genetic deletion of LRRK2 prevented the ΔpH-induced vacuolation, implying that the kinase is required to modulate vesicular traffic. We propose that by dictating the state of activation of LRRK2 and in turn that of Rab GTPases, the development of a progressive luminal acidification serves as a timing device to control endocytic maturation.

## INTRODUCTION

Numerous ligands and their cognate plasmalemmal receptors, as well as solutes present in the extracellular fluid phase are internalized and sorted for recycling, transport or eventual degradation as they travel along the various compartments of the endocytic pathway. The pH of these compartments is not homogeneous, becoming increasingly acidic as early endosomes progress to late endosomes and ultimately lysosomes ^1,2^. Thus, the luminal fluid that initially has a pH similar to the trapped extracellular milieu (i.e. 7.2-7.4) can reach ≈4.5 in terminal lysosomes.

The characteristic pH of each individual stage of the endocytic pathway is dictated by the balance between the rates of inward proton pumping and passive outward “leakage” of proton equivalents ^3^. Proton pumping, which is mediated by vacuolar H^+^-ATPases (V-ATPases) ^4–6^, is in turn determined by their density and inherent transport rate. Because V-ATPases are electrogenic, proton pumping is sensitive to the transmembrane potential, a function of the ionic permeability of the membrane. The more pronounced acidification attained by late endosomes and lysosomes is attributed to their greater abundance of V-ATPases, higher counterion conductance and reduced rates of proton leakage ^7,8^.

The role of the varying pH along the endocytic pathway in the dissociation and recycling of ligands and receptors has been well established ^9–12^, and the contribution of acidification to cargo degradation is also widely acknowledged ^13–17^. In addition, a variety of studies indicate that luminal acidification is essential for the normal traffic of membranes between the compartments of the endocytic pathway. Perturbation of the existing organellar acidification, whether by inhibiting the V-ATPases or by counteracting their activity using weak bases or protonophores has been shown to impair the formation of endocytic carrier vesicles ^18–21^, endosome tubulation and recycling ^18,22–28^, and the transition between early and late compartments ^17,29–32^.

Despite the abundance of evidence accumulated over decades, and while some hypotheses have been put forward (see Discussion), no clear consensus has emerged regarding the mechanism by which luminal pH controls endocytic traffic. We revisited this important unanswered question using macrophages as an experimental model. Endocytosis is uniquely active in macrophages, which internalize the equivalent of their entire cell surface every ≈20 min ^33^, facilitating the detection of events that impair traffic. Our data indicate that the luminal pH of endosomes dictates the state of activation of Rab GTPases, a process regulated by LRRK2, a kinase implicated in the etiology of Parkinson’s disease. In this manner, the pH influences the transition between the various stages of the endocytic pathway.

## RESULTS

### Dissipation of the transmembrane pH gradient (ΔpH) induces rapid vacuolation that is dependent on Rab5

To study the effects of luminal pH on endocytic traffic RAW 264.7 macrophages (hereafter called RAW) were treated with a combination of concanamycin A, a potent and specific inhibitor of proton (H^+^) pumping by the V-ATPase, and nigericin, a K^+^/H^+^ exchanger. By combining these two agents we ensured that the transmembrane pH would be inhibited rapidly, without incurring marked osmotic changes or accelerating ATP consumption by uncoupled proton pumps. Distinct morphological changes were apparent by differential interference contrast (DIC) microscopy within 30 min of acute ΔpH dissipation (Fig. 1a). Multiple large (∼ 2 µm) vacuoles appeared in the cells treated with concanamycin/nigericin (Fig. 1a, right), which were rare in untreated macrophages (Fig. 1a, left). The formation of these vacuoles was also readily apparent by transmission electron microscopy (TEM), which additionally showed the presence of intravacuolar vesicles (Fig. 1b).

**Figure 1.**
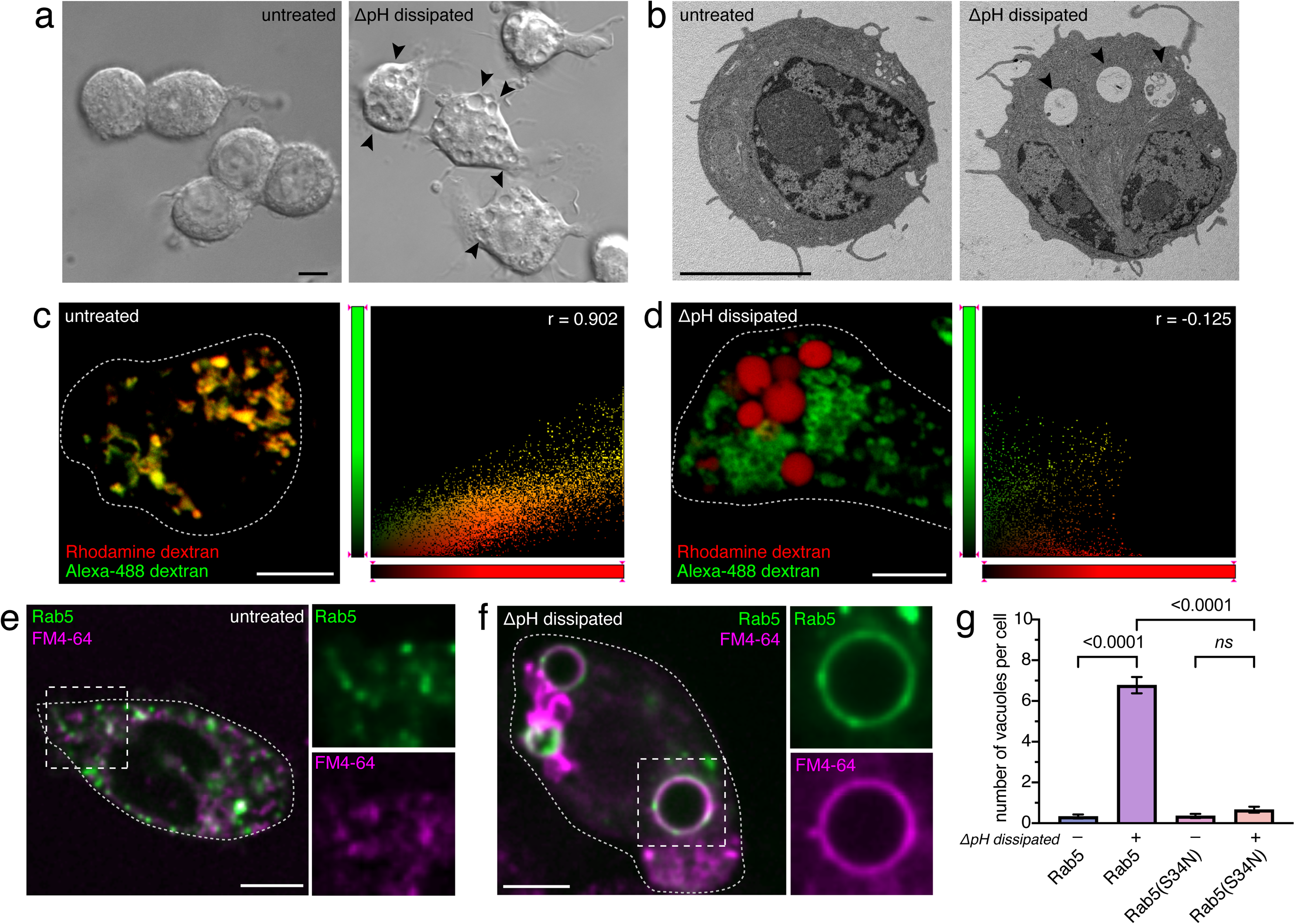
ΔpH dissipation induces rapid vacuolation that is dependent on Rab5. For this and subsequent figures, luminal acidification was dissipated by 30 min treatment with 250 nM concanamycin A and 5 µg·mL^−1^ nigericin at 37°C in isotonic K^+^ medium. **a.** Untreated and ΔpH-dissipated RAW 264.7 macrophages (hereafter called RAW) visualized by differential interference contrast (DIC) microscopy. Arrowheads point to vacuoles formed in ΔpH-dissipated cells. **b.** Untreated and ΔpH-dissipated RAW macrophages visualized by transmission electron microscopy (TEM). Arrowheads point to vacuoles formed in ΔpH-dissipated cells. **c.** RAW cells were pulsed with Alexa Fluor 488-conjugated dextran (green) for 16 h followed by a 1 h chase to label primarily lysosomes. Rhodamine-conjugated dextran (red) was then pulsed for 30 min, removed from the medium and chased for an additional 30 min. Cell outline indicated by dotted line. Colocalization scatter plot is shown at right; r = Pearson’s coefficient. **d.** RAW cells were treated as in **c.**, except that cells were ΔpH dissipated during the pulse with rhodamine-conjugated dextran as well as during the chase. Scatter plot is shown at right; r = Pearson’s coefficient. **e.** Untreated RAW cells pre-labeled with internalized FM4-64 (magenta) and expressing wildtype Rab5 (green). Side panels show individual Rab5 and FM4-64 channels at 1.8× magnification. **f.** ΔpH-dissipated RAW cells labeled with FM4-64 and Rab5 (green) as in **e**. Side panels show individual channels at 1.6× magnification. **g.** Number of FM4-64^+^ vacuoles quantitated in untreated and ΔpH-dissipated RAW cells expressing either wildtype Rab5 or Rab5(S34N). For each condition, 3 independent experiments were quantified, with ≥ 10 cells per replicate. Data are means ± SEM. *p* calculated using ordinary one-way ANOVA. All images are representative of ≥ 30 fields from ≥ 3 experiments of each type. All scale bars: 5 µm.

Vacuolation coincided with and likely resulted from impaired endocytic traffic between the early and late compartments (Fig. 1c-d). This was apparent when comparing the fate of endocytic markers in untreated cells and in cells where the ΔpH was dissipated. To this end, late endosomes/lysosomes were initially labeled with green (Alexa 488-labeled) dextran using a standard pulse-chase protocol. In otherwise untreated cells, a subsequent pulse with red (rhodamine-labeled) dextran resulted in extensive mixing of the two markers within 30 min (Fig.1c; colocalization scattergram at right), indicative of fusion of early with late compartments. By contrast, little colocalization was observed in cells where pH had been neutralized by concanamycin/nigericin immediately before delivering the second marker (Fig. 1d, colocalization scattergram at right). Similar vacuolation and traffic arrest were observed when concanamycin was combined with ammonium to induce rapid pH dissipation (not illustrated); for simplicity, only concanamycin/nigericin was used hereafter.

To confirm that the large vacuoles generated by pH-dissipated cells were of endocytic nature, the cells were pre-loaded with FM4-64, an amphiphilic dye that inserts into the plasma membrane and is rapidly internalized. Because it is membrane impermeant and binds reversibly, plasmalemmal-associated FM4-64 can be readily washed from the cell surface, revealing exclusively endocytic organelles. As shown in Fig. 1e, while the endocytic tracer labels primarily small vesicles and tubules in otherwise untreated cells, it demarcates the large vacuoles observed by DIC in ΔpH-dissipated cells (Fig. 1f). Of note, a substantial fraction of the vesicles and vacuoles are also labeled by Rab5, that was labeled with GFP and expressed ectopically (Figs. 1e-f). Whereas multiple Rab5-positive large vacuoles were observed in cells exposed to concanamycin/nigericin (6.78 ± 0.40 vacuoles per cell), they were infrequent in macrophages (0.32 ± 0.09 per cell; Fig. 1e). The alterations induced by ΔpH dissipation were not restricted to macrophages, though they were less pronounced in cells with lower endocytic activity. The distribution of Rab5 was also affected in HeLa cells when treated with concanamycin/nigericin (Extended Data Fig. 1). After ΔpH dissipation, the size and fluorescence intensity of the punctate Rab5-positive structures increased markedly, while the diffuse cytoplasmic Rab5 fluorescence decreased concomitantly (*cf.* Extended Data Figs. 1a and b). As a result, the ratio of punctate to cytoplasmic Rab5 fluorescence nearly tripled (Extended Data Fig. 1c).

Because the Rab5 GTPase is proposed to be a master regulator of endosome biogenesis and traffic, we considered whether its activity was required for the observed vacuolar enlargement. To this end, we assessed the effects of ΔpH dissipation in cells expressing a dominant-negative form of Rab5 (Rab5(S34N)). As illustrated in Fig. 1g, Rab5(S34N) virtually eliminated the formation of large vacuoles in cells treated with concanamycin/nigericin. This indicated that formation of the vacuoles was dependent on Rab5 function.

### ΔpH dissipation results in the hyperactivation of Rab GTPases

Interestingly, the large vacuoles induced by ΔpH dissipation are reminiscent of the enlarged endocytic structures observed in cells that express constitutively-active Rab5 ^23,34^ (and our unpublished observations in macrophages). Therefore, we sought to determine the activation state of Rab5 in ΔpH-dissipated cells. A biosensor for GTP-bound Rab5, the Rabaptin 5-binding domain (R5BD) that selectively binds the active form of the GTPase, was co-expressed in cells along with tagged wild-type Rab5. In untreated macrophages, the probe localized to the cytosol and in punctate Rab5-positive structures that were likely early endosomes bearing active Rab5 (Fig. 2a). In ΔpH dissipated cells, the probe bound to the large vacuoles (Fig. 2b). This phenomenon was also observed in HeLa cells, where the R5BD probe bound more strongly to the enlarged Rab5-positive vesicles of ΔpH-dissipated cells (Extended Data Fig. 2a-b).

**Figure 2.**
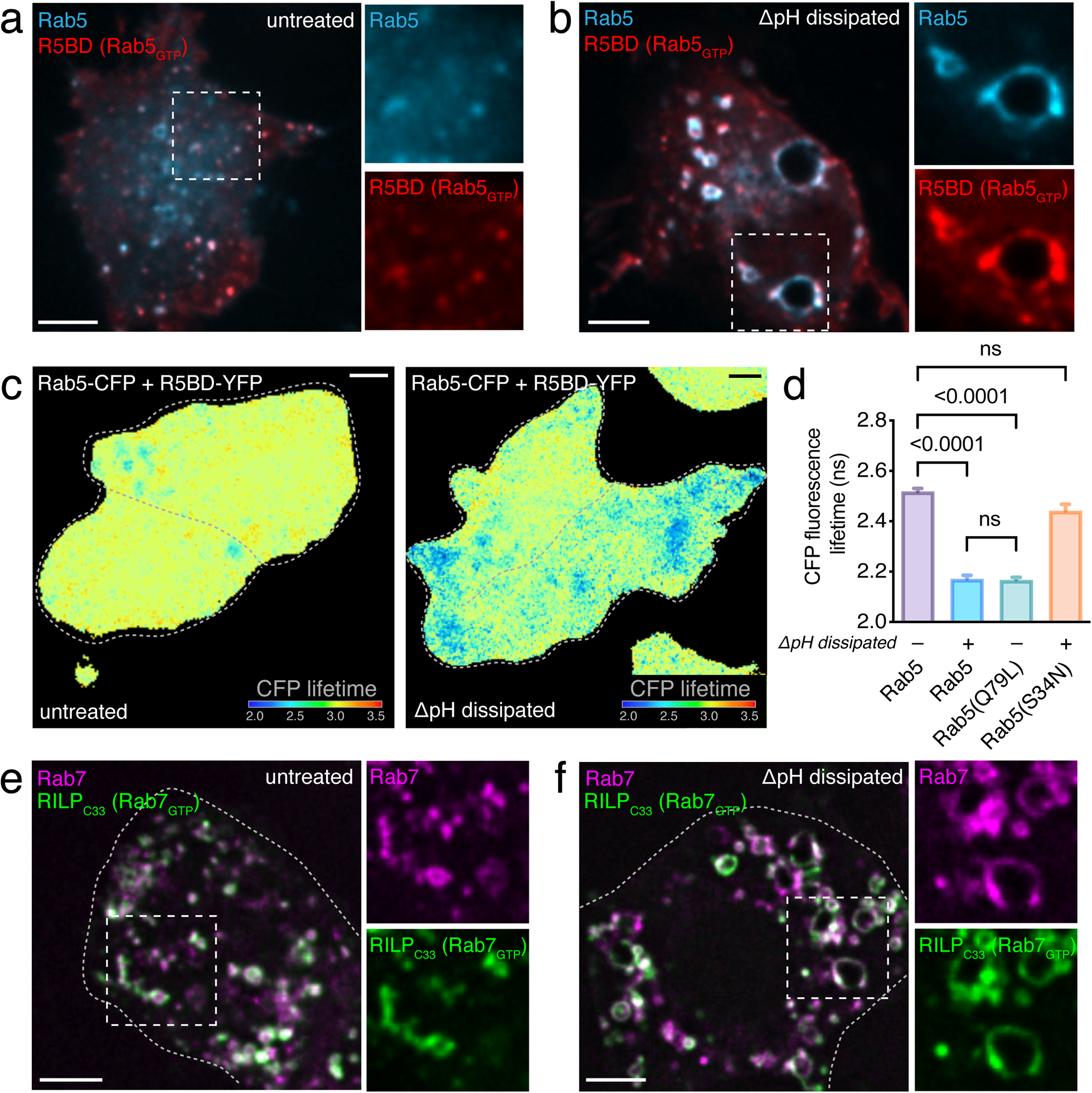
ΔpH dissipation results in the hyperactivation of Rab GTPases on enlarged endocytic vacuoles. **a.** Untreated RAW cells expressing wildtype Rab5 (cyan) and R5BD (red), a biosensor for GTP-bound Rab5. Side panels show individual Rab5 and R5BD channels at 1.9× magnification. **b.** ΔpH-dissipated RAW cells expressing Rab5 (cyan) and R5BD (red). Side panels show individual Rab5 and R5BD channels at 1.7× magnification. **c.** Pseudocolored fluorescence lifetime (FLIM) images of untreated (left) an ΔpH-dissipated RAW cells (right) co-expressing CFP-Rab5 and YFP-R5BD. CFP lifetime is pseudocolored in a rainbow LUT corresponding to lifetime values from 2 to 3.5 ns. **d.** CFP fluorescence lifetimes measured for untreated or ΔpH-dissipated cells co-expressing CFP-Rab5, CFP-Rab5(Q79L) or CFP-Rab5(S34N) and YFP-R5BD. For each condition, 3 independent experiments were quantified, with ≥ 20 cells per replicate. Data are means ± SEM. *p* calculated using ordinary one-way ANOVA. **e.** Untreated RAW cells expressing Rab7 (magenta) and RILP_C33_ (green), a biosensor for GTP-bound Rab7. Side panels show individual Rab7 and RILP_C33_ channels at 1.5× magnification. **f.** ΔpH-dissipated RAW cells expressing Rab7 (magenta) and RILP_C33_ (green). Side panels show individual Rab7 and RILP_C33_ channels at 1.6× magnification. All images are representative of ≥ 30 fields from ≥ 3 experiments of each type. Cell outlines are indicated by dotted lines. All scale bars: 5 µm.

The association between R5BD and Rab5 in macrophages was quantified by Förster resonance energy transfer (FRET), measured by fluorescence lifetime imaging microscopy (FLIM). Measurement of FRET by FLIM is advantageous, as it is independent of the absolute concentration of the fluorophore and unaffected by photobleaching ^35^. In FLIM-FRET, the distance-dependent energy transfer between donor and acceptor molecules results in a measurable shortening of donor fluorescence lifetime; FRET efficiency can thus be calculated by comparing the lifetime of the donor fluorescence when expressed by itself or jointly with the acceptor. In otherwise untreated macrophages expressing CFP-Rab5 (donor) and YFP-R5BD (acceptor), the lifetime of the CFP associated with punctate, endosome-like structures averaged 2.45 ± 0.01 ns (Fig. 2c, left, and d), equivalent to a FRET efficiency of 4.9%. Notably, CFP fluorescence lifetime was markedly depressed on the large vacuole-like structures formed in ΔpH dissipated cells (2.17 ± 0.01 ns; Fig. 2c, right, and d), corresponding to a FRET efficiency of 13.8%. This lifetime decrease was similar to that seen in cells expressing CFP-tagged constitutively-active Rab5 (Rab5(Q79L) with YFP-R5BD (Fig. 2d; FRET efficiency = 15.7%). In contrast, pH dissipation had little effect on the fluorescence lifetime of cells expressing the dominant negative CFP-Rab5(S34N) along with YFP-R5BD, yielding a low FRET efficiency (1.5%; Fig. 2d). These data strongly suggested that dissipation of the endosomal pH gradient resulted in hyperactivation of Rab5, thereby promoting the formation of large vacuoles.

Remarkably, the GTPase hyperactivation was not restricted to the Rab5 compartment. Enlarged Rab7-positive vacuoles were also observed in cells where ΔpH had been dissipated (Fig. 2f), that were distinctly different from the small Rab7-containing vesicles and tubules found in untreated cells (Fig. 2e). The enlarged vacuoles contained GTP-bound Rab7, as indicated by their ability to bind RILP_C33_, a biosensor for active Rab7 ^36^. Therefore, dissipation of the pH gradient resulted in hyperactivation of at least two Rab GTPases, suggesting a common underlying mechanism.

### Hyperactivation of Rab GTPases during ΔpH dissipation is the result of inhibited GAP activity

The mechanism underlying the apparent Rab hyperactivation was probed next. The state of activation of Rab GTPases is determined by the balance between the exchange of GDP for GTP and the hydrolysis of GTP. These functions are mediated by GEF and GAP proteins, respectively. Therefore, in principle, hyperactivation of Rab5 or Rab7 could be caused by stimulating GEF or by inhibiting GAP activity. Fluorescence recovery after photobleaching (FRAP) of CFP/GFP-tagged Rabs was used to discern between these alternatives. Recovery of fluorescence following bleaching of an entire, individual vesicle or vacuole can only occur by dissociation of the bound, photobleached Rab upon conversion to its GDP-bound form, followed by binding of cytosolic fluorescent Rab in the GTP-bound form. Rab hyperactivation due to accelerated nucleotide exchange (GEF activity) would be expected to result in faster recovery after photobleaching, while slower or incomplete recovery would indicate GAP inhibition.

The validity of this approach was ascertained by comparing the rates of recovery of wildtype and constitutively active Rab5 (Rab5(Q79L). As illustrated in Fig. 3a, whereas recovery of wildtype Rab5 in untreated cells was rapid (t_1/2_ = 17.1 s) and extensive (75.6%), the active form recovered much more slowly (t_1/2_ = 26.6 s) and incompletely (fractional recovery 18.9%) during the 90 s observation period. Similar results were obtained comparing the recovery of wildtype and constitutively active Rab7 (Rab7(Q67L); t_1/2_ = 17.3 s vs. 32.3 s and recovery = 83.9% vs. 33.8%; Fig 3b). Importantly, in cells where ΔpH had been dissipated both Rab5 and Rab7 recovered more slowly (t_1/2_ = 27.8 s and 22.9 s), respectively, and not as completely (61.6% and 40.8%) as they did in untreated cells. These observations were replicated in HeLa cells (Extended Data Fig. 2c), where the enlargement of the compartments induced by altering the pH is not as pronounced, facilitating their selective bleaching. These results are indicative of slower Rab dissociation and can be most readily explained by a pH-dependent reduction in the rate of conversion from the active (GTP- and membrane-bound) form of the Rab to the dissociable (GDP-bound) form.

**Figure 3.**
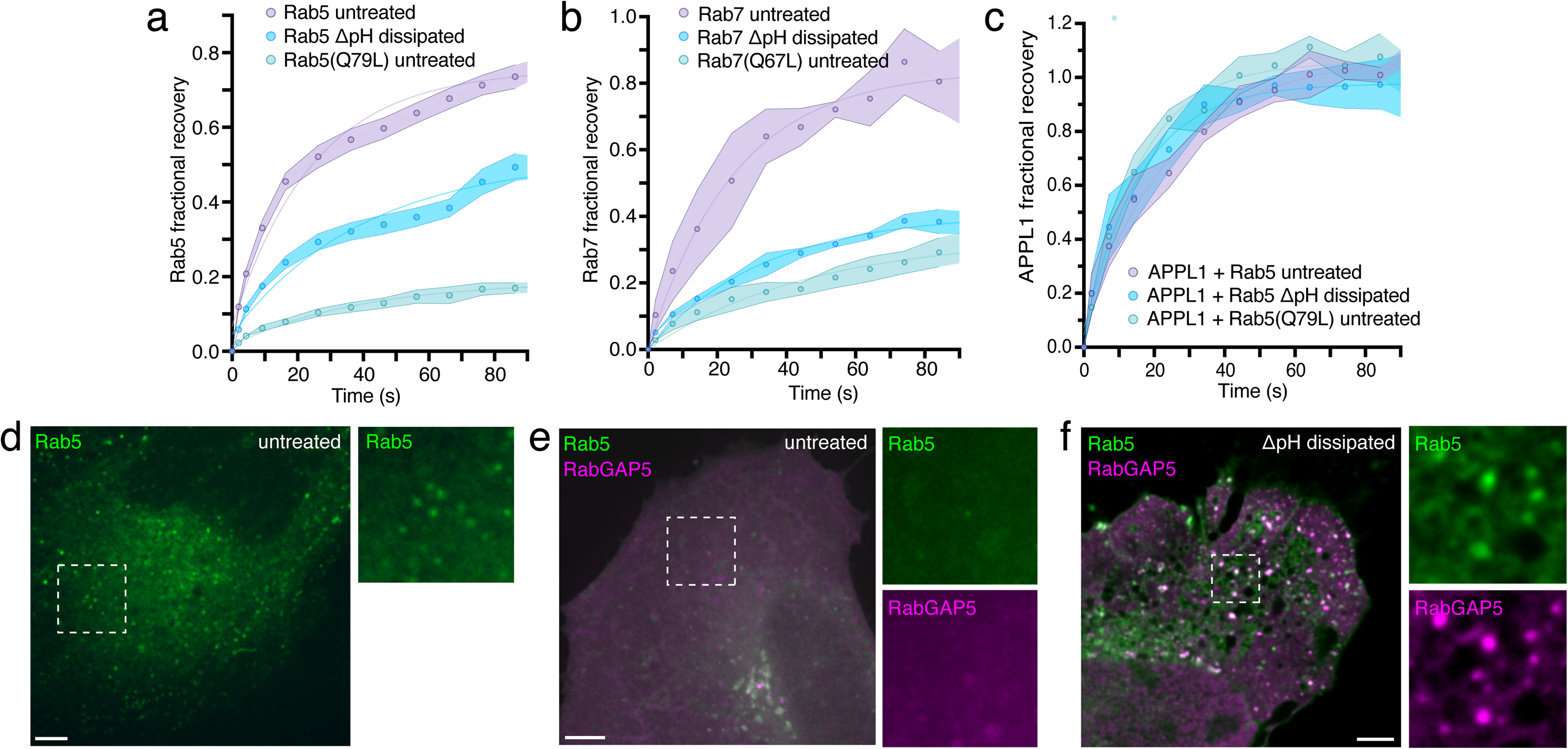
Activation of Rab GTPases upon ΔpH dissipation is the result of inhibited GAP activity. **a.** Entire Rab5-positive vesicles were photobleached in untreated or ΔpH-dissipated RAW cells expressing CFP-Rab5 or CFP-Rab5(Q79L), and the vesicle Rab5 fluorescence recovery monitored. For this and all FRAP curves, average vesicle fluorescence before photobleaching was defined as 1.0, and fluorescence recovery normalized to that of an unbleached vesicle to correct for bleaching incurred during the acquisition of images during the recovery phase. For each condition, 3 independent experiments were quantified, with ≥ 10 cells per replicate. Data are means ± SEM. **b.** Entire Rab7-positive vesicles were photobleached in untreated or ΔpH-dissipated RAW cells expressing GFP-Rab7 or GFP-Rab7(Q67L), and the vesicle Rab7 fluorescence recovery monitored. For each condition, 3 independent experiments were quantified, with ≥ 10 cells per replicate. Data are means ± SEM. **c.** Entire Rab5-positive vesicles were photobleached in untreated or ΔpH-dissipated RAW cells co-expressing CFP-Rab5 or CFP-Rab5(Q79L) and GFP-APPL1, and the APPL1 fluorescence recovery of individual vesicles monitored. For each condition, 3 independent experiments were quantified, with ≥ 10 cells per replicate. Data are means ± SEM. **d.** Untreated HeLa cells expressing wildtype Rab5 (green). Side-panel shows Rab5 channel at 2.3× magnification. **e.** Untreated HeLa cells expressing Rab5 (green) and RabGAP5 (magenta). Side panels show individual Rab5 and RabGAP5 channels at 2.3× magnification. **f.** ΔpH-dissipated HeLa cells expressing Rab5 (green) and RabGAP5 (magenta). Side panels show individual Rab5 and RabGAP5 channels at 3.3× magnification. All images are representative of ≥ 30 fields from ≥ 3 experiments of each type. All scale bars: 5 µm.

A reduced rate of GTP hydrolysis may be attributable to inhibition of GAP activity or to failure of the GAP to gain access to the GTP-bound Rab. The latter could result from more persistent association with effectors. This possibility was analyzed measuring the rate of dissociation of APPL1, a Rab5 effector that associates preferentially with the active form of the GTPase. As shown in Fig. 3c, the recovery of fluorescence of membrane-bound GFP-APPL1 was rapid and unaffected by dissipation of the ΔpH and was similar in cells expressing wildtype and constitutively active Rab5. Therefore, persistent association of the effector cannot account for the reduced recovery of Rab5 fluorescence in cells where the ΔpH was dissipated. Inhibition of GAP activity is therefore a more likely explanation.

We also considered whether the persistent activation of Rab5 could be secondary to a block in the maturation between early and late endosomal compartments. However, co-expression of Rab5 with dominant-negative or constitutively-active Rab7 constructs (Rab7(T22N) or Rab7(Q67L, respectively) did not replicate or alleviate the poor recovery of Rab5 induced by ΔpH dissipation (Extended Data Fig 3b-c, compared to panel a, where wild type Rab7 is co-expressed with Rab5). These observations imply that the activation of Rab5 by manipulating the pH is independent of, and likely the cause of the impaired transition between the early and late stages (Fig. 1f-g).

Because reduced GTPase activity seemed the most likely explanation, we proceeded to analyze whether pH dissipation blocked the ability of Rab GAPs to localize to the endosomes. Two different GAPs, which have been shown to target Rab5, were tested. When overexpressed ectopically RabGAP5 displaced Rab5 from endosomes, confirming its effectiveness as a GAP for this GTPase (Fig 3d-e). These experiments were performed in HeLa cells, where (over)expression of multiple plasmids can be accomplished much more readily than in macrophages. In contrast, following dissipation of the ΔpH, wildtype Rab5 remained associated with endomembranes despite co-expression of RabGAP5 (Fig 3f). Importantly, RabGAP5 bound effectively to the endosomes of ΔpH-dissipated cells, implying that the latter displayed active (GTP-bound) Rab5 and that inability to bind the GAP was not responsible for the increased Rab activity. That RabGAP5 contributes to the endogenous GAP activity of HeLa cells was demonstrated using siRNA: silencing its expression caused enlargement of the Rab5-positive compartments (Extended Data Fig. 4c), resembling the effects of pH dissipation, and reduced the rate and extent of Rab5 fluorescence recovery in FRAP experiments (Extended Data Fig. 4d). Similar results were obtained using a second, less well characterized Rab5 GAP, namely TBC1D2B, which displaced Rab5 from endosomes in otherwise untreated cells, but failed to do so following elimination of the pH gradient (Extended Data Fig. 4a). Silencing of TBC1D2B also enlarged the Rab5 compartment and reduced the dissociation of the GTPase as visualized by FRAP, effects that were even more pronounced when RabGAP5 was simultaneously silenced (Extended Data Fig. 4d). Jointly, these data are consistent with the notion that neutralization of pH inhibits the function of GAPs, without impairing their ability to bind to active Rab5.

### LRRK2 localization and activity are pH-dependent

Because at least two different Rab proteins, Rab5 and Rab7, were hyperactivated in a pH-dependent manner, a common underlying mechanism was considered likely. Recently, a Parkinson’s disease-related kinase, leucine-rich repeat kinase 2 (LRRK2), was shown to phosphorylate a subset of endocytic Rab GTPases ^37^, rendering them insensitive to GAP-mediated GTP hydrolysis ^37–40^. While the activation of LRRK2 has been linked to the autophosphorylation of a key serine (S1292) ^41–43^, the exact cellular signal triggering this activation has not been identified. We therefore sought to investigate whether the vacuolation phenotype seen during ΔpH dissipation was linked to LRRK2.

LRRK2 is a very large protein (286 kDa) and the unwieldy plasmids encoding it are difficult to introduce and express in most cells. To overcome this limitation, we used AD293 cells. Fluorescently-tagged LRRK2 was expressed in these cells and its localization analyzed before and after neutralization of the pH. As in previous reports, LRRK2 was found to localize mainly to the cytosol of untreated cells, with little accumulation at discrete sites (9.0 ± 1.4 puncta per cell; see Fig. 4a and d). However, upon pH dissipation, the number and brightness of LRRK2 puncta increased markedly (37.4 ± 4.6 per cell, *p* <0.0001; see Fig. 4b and e). The LRRK2 puncta appeared to localize in the immediate vicinity of the enlarged Rab5 and Rab7 endosomes that appeared after ΔpH dissipation (Fig. 4b and e). The significance of the coupling between LRRK2 and Rab domains was computed using Statistical Object Distance Analysis (SODA). SODA utilizes the morphology and distance separating coupled structures to provide a statistical map of intracellular molecular assemblies at a population level, which is independent of point spread function and is robust to noise ^44^. SODA coupling analyses revealed that ΔpH dissipation increased the coupling of Rab5- or Rab7-bearing membranes to LRRK2 puncta by 9.1-fold and 3.6-fold, respectively (Fig. 4c and f).

**Figure 4.**
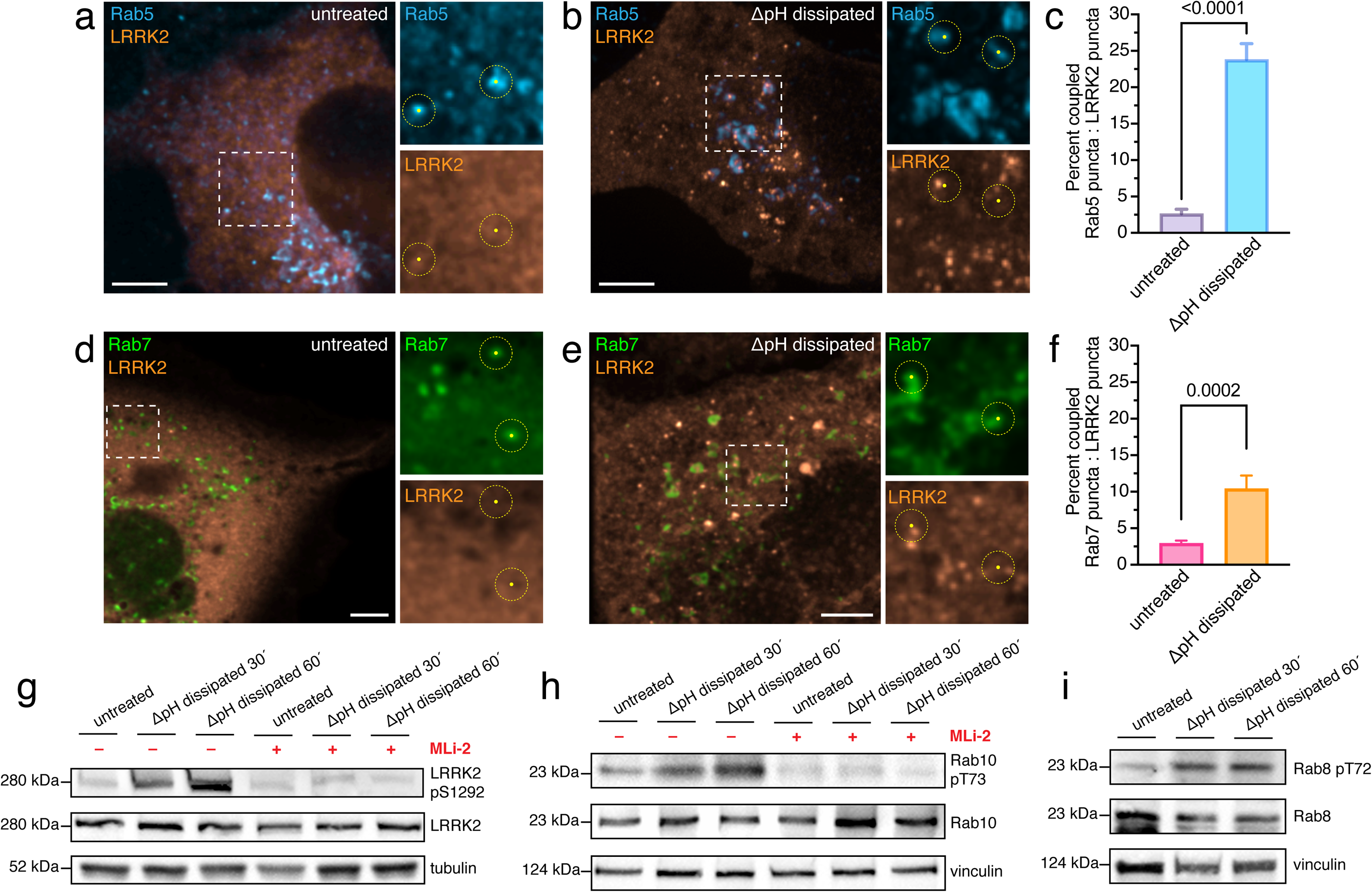
LRRK2 localization and activity is pH-dependent and leads to the phosphorylation of endocytic Rab GTPases. **a.** Untreated AD293 cells expressing Rab5 (cyan) and LRRK2 (orange). Side panels show individual Rab5 and LRRK2 channels at 2.0× magnification. For this and subsequent panels, the immediate are (∼500 nm radius) surrounding identified Rab^+^ vesicles (yellow dots) utilized in SODA coupling analysis is represented by dotted yellow circles. **b.** ΔpH-dissipated AD293 cells expressing Rab5 (cyan) and LRRK2 (orange). Side panels show individual Rab5 and LRRK2 channels at 1.9× magnification. **c**. Percent of Rab5-positive puncta coupled to LRRK2-positive puncta in untreated and ΔpH-dissipated AD293 cells. Percentages calculated from SODA coupling analysis using Icy software (http://icy.bioimageanalysis.org/). For each condition, 3 independent experiments were quantified, with ≥ 10 cells per replicate. Data are means ± SEM. *p* calculated using unpaired, 2-tailed student’s t-test. **d.** Untreated AD293 cells expressing Rab7 (green) and LRRK2 (orange). Side panels show individual Rab7 and LRRK2 channels at 2.7× magnification. **e.** ΔpH-dissipated AD293 cells expressing Rab7 (green) and LRRK2 (orange). Side panels show individual Rab5 and LRRK2 channels at 2.4× magnification. **f**. Percent of statistically coupled Rab7-positive puncta to LRRK2-positive puncta in untreated and ΔpH-dissipated AD293 cells. Percentages calculated from SODA coupling analysis. For each condition, 3 independent experiments were quantified, with ≥ 10 cells per replicate. Data are means ± SEM. *p* calculated using unpaired, 2-tailed student’s t-test. **g**. LRRK2 pS1292 immunoblot of lysates prepared from RAW cells that were untreated or ΔpH-dissipated for the indicated times, in the absence and presence of 50 nM MLi-2, as indicated. Total LRRK2 and tubulin immunoblots of identical aliquots shown as loading controls. **h**. Rab10 pT73 immunoblot of lysates prepared from RAW cells that were untreated or ΔpH-dissipated for indicated times, in the absence and presence of 50 nM MLi-2, as noted. Total Rab10 and vinculin immunoblots shown as loading controls. **i**. Rab8 pT72 immunoblot of lysates prepared from RAW cells that were untreated or ΔpH-dissipated for indicated times, in the absence and presence of MLi-2. Rab8 pT72 blot is shown alongside total Rab8 and vinculin loading controls. Blots in **g**, **h** and **i** are representative of 3 independent experiments of each kind. All images are representative of ≥ 30 fields from ≥ 3 experiments of each type. All scale bars: 5 µm.

LRRK2 has been shown to phosphorylate Rab proteins in response to seemingly disparate conditions such as lysosomal rupture, cell exposure to weak bases, and incubation with lysosomotropic agents ^45–48^. We hypothesized that the unifying feature of all these conditions is the resultant dissipation of the endosome/lysosome transmembrane pH gradient. Therefore, we tested whether LRRK2 was activated in macrophages using our ΔpH dissipation protocol. Indeed, exposure to concanamycin/nigericin for 30 or 60 min resulted in robust autophosphorylation of LRRK2 at S1292 that was sensitive to pretreatment with the LRRK2-specific inhibitor MLi-2 ^49,50^ (Fig. 4g). As there are currently no reliable antibodies to detect phosphorylated forms of Rab5 or Rab7, we used available phospho-specific antibodies to test the phosphorylation of Rab10 and Rab8 ^51^, which are well-established LRRK2 substrates. Both were found to become unequivocally phosphorylated in response to ΔpH dissipation, in a LRRK2-dependent manner (Fig. 4h-i). Of interest, the membrane compartments bearing these GTPases were also enlarged by ΔpH dissipation, though not as markedly as the Rab5 compartment (Extended Data Fig. 5a-b). Jointly, these data suggest that the activating signal for LRRK2 is, in fact, disruption of the endo-lysosomal pH gradient, which induces LRRK2 coupling to endomembranes. Following its recruitment and activation LRRK2 phosphorylates membrane-bound Rab GTPases, thereby locking them in GTP-bound, hyperactive states insensitive to deactivation by GAPs.

### LRRK2 is activated in the absence of membrane damage and is required for the ΔpH-induced vacuolation

Previous reports describe LRRK2 activation as part of the cellular response to lysosomal membrane damage ^46,48^. Therefore, we considered the possibility that membrane rupture was occurring under the conditions we used to dissipate the pH. Whether membrane damage occurred during concanamycin/nigericin-mediated ΔpH dissipation was tested by preloading the endocytic pathway with sulforhodamine B, a small dye with a Stokes radius < 1 nm. This dye enabled visualization of membrane rupture: exposure of cells to LLOMe, an agent used previously to induce LRRK2 activation ^46,48^, caused release of the vesicular sulforhodamine B to the cytosol (Fig. 5a, right 2 panels). In sharp contrast, the enlarged Rab5- or Rab7-positive vacuoles formed in cells treated with concanamycin/nigericin retained the dye within their lumen. This suggested that membrane damage was not required for LRRK2 activation. Therefore, it appears more likely that the dissipation of luminal pH that inevitably accompanies lysosome rupture –rather than membrane rupture itself– was the cause of the observed activation of LRRK2 in cells treated with LLOMe or other agents of membrane damage ^45–48^.

**Figure 5.**
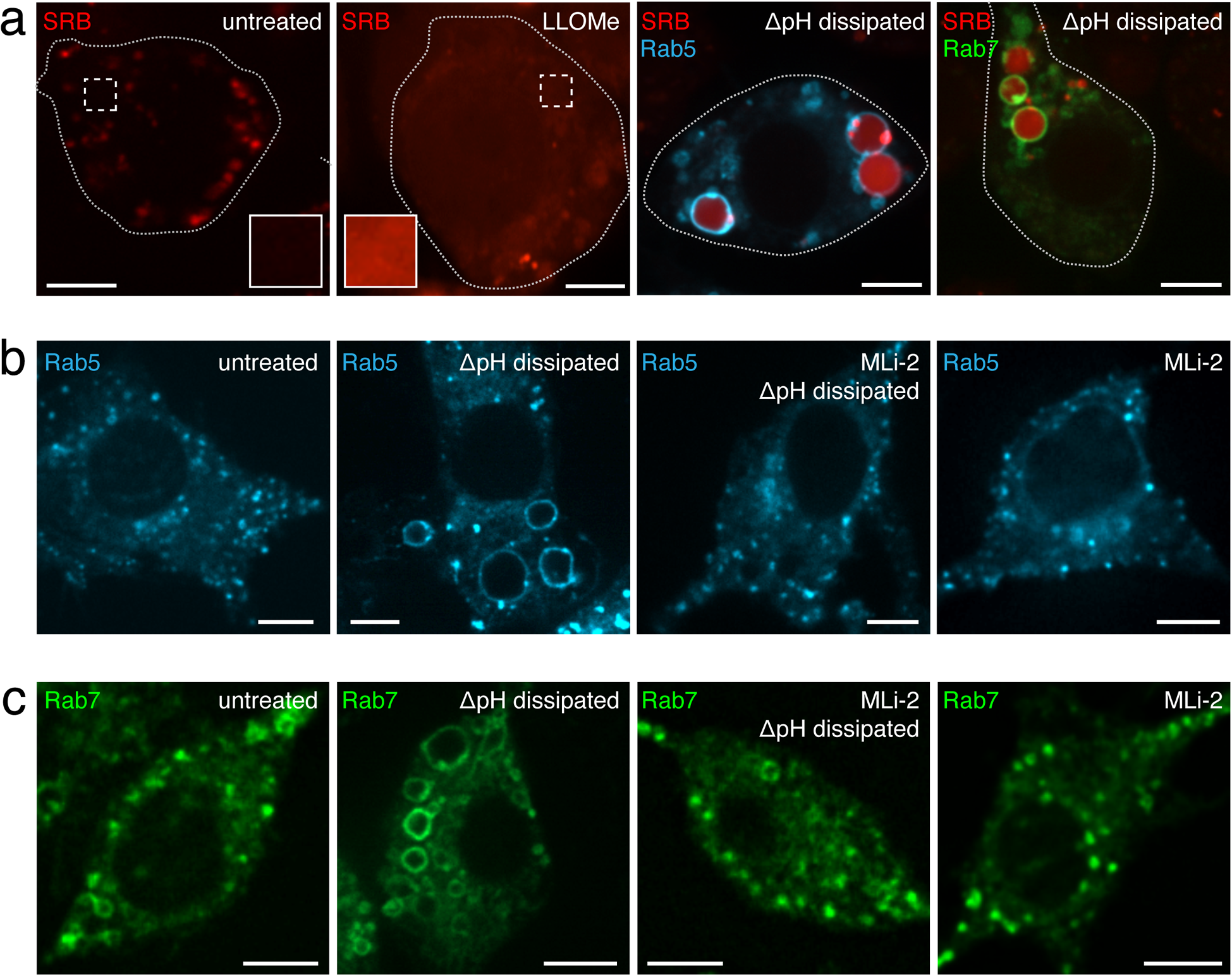
LRRK2 is activated in the absence of membrane damage and is required for the vacuolation caused by ΔpH dissipation. **a**. Comparison of sulforhodamine B (SRB; red) retention within endolysosomes of untreated, LLOMe-treated, and ΔpH-dissipated RAW cells expressing Rab5 (cyan) or Rab7 (green). Insets in untreated and LLOMe-treated panels show an area of cytoplasm with intensity enhanced 2× and magnified at 2.3× to better visualize leakage of SRB. **b.** Effect of MLi-2 treatment on the vacuolation of Rab5-positive membranes (cyan) induced by ΔpH dissipation in RAW cells. **c.** Effect of MLi-2 treatment on the vacuolation of Rab7-positive membranes (green) induced by ΔpH dissipation. All images are representative of ≥ 30 fields from ≥ 3 experiments of each type. All scale bars: 5 µm.

Next, it was important to establish whether the activation of LRRK2 was responsible for the enlargement of the endocytic membranes, or was merely an associated event. To this end, cells were pretreated with MLi-2 prior to and during ΔpH dissipation. Remarkably, MLi-2 –which had no discernible effects on otherwise untreated cells (Figs. 5b and c, rightmost panels)– completely blocked the formation of Rab5- or Rab7-positive vacuoles in response to concanamycin/nigericin (*cf.* middle panels in Figs. 5b and c). Inhibition of LRRK2 also blocked the pH-induced enlargement of the Rab10 and Rab8 compartments (Extended Data Fig. 5a-b). These data suggest that LRRK2 activation is causally related to the vacuolation of endocytic compartments.

Because pharmacologic agents can potentially affect unintended targets, we sought additional means to test the involvement of LRRK2 in the pH-dependent alterations in endocytic traffic. With this aim we tested the effects of ΔpH dissipation in RAW macrophages where the *LRRK2* gene had been deleted. Immunoblotting confirmed the absence of LRRK2 protein in the LRRK2^−/−^ RAW cells (Fig. 6a). As anticipated, Rab10 or Rab8 phosphorylation in response to concanamycin/nigericin or to exposure to chloroquine –a weak base that effectively neutralizes luminal pH– was absent in the LRRK2^−/−^ cells (Fig. 6b-d). Most importantly, vacuolation failed to occur when ΔpH was dissipated in the knockout cells (Fig. 6d-e). The Rab10 and Rab8 compartments similarly failed to expand in the LRRK2^−/−^ cells (Extended Data Fig 6a-b).

**Figure 6.**
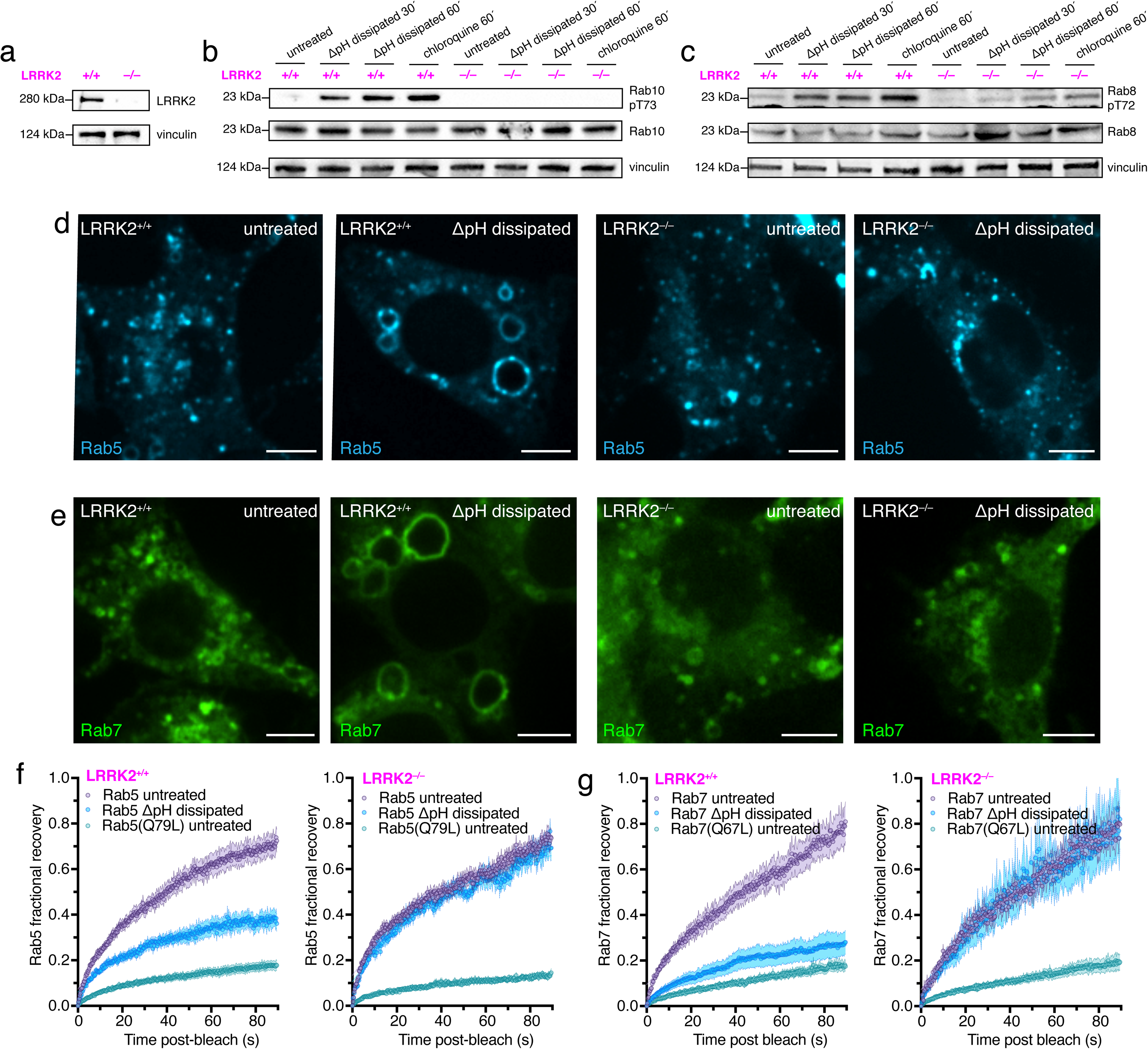
LRRK2 is required for the pH-induced refractoriness to Rab-GAP activity. **a**. LRRK2 immunoblot of LRRK2^+/+^ and LRRK2^−/−^ RAW cell lysates. Vinculin immunoblot is shown as loading control. **b.** Rab10 pT73 immunoblot of lysates prepared from LRRK2^+/+^ or LRRK2^−/–-^RAW cells that were untreated or ΔpH-dissipated for indicated times. Total Rab10 and vinculin immunoblots of identical aliquots shown as loading controls. **c**. Rab8 pT72 immunoblot of lysates prepared from LRRK2^+/+^ or LRRK2^−/−^ RAW cells that were untreated or ΔpH-dissipated for indicated times. Total Rab8 and vinculin immunoblots shown as loading controls. Blots in **a**, **b** and **c** are representative of 3 independent experiments of each kind. **d.** Effect of LRRK2 knock out on the vacuolation of Rab5-positive membranes (cyan) induced by the ΔpH dissipation in RAW cells. **e.** Effect of LRRK2 knock out on the vacuolation of Rab7-positive membranes (green) induced by the ΔpH dissipation. All images in **d** and **e** are representative of ≥ 30 fields from ≥ 3 experiments of each type. All scale bars: 5 µm. **f.** Comparison of Rab5 FRAP in LRRK2^+/+^ (left) versus LRRK2^−/−^ RAW cells (right). FRAP was measured as described in untreated or ΔpH-dissipated LRRK2^+/+^ or LRRK2^−/−^ RAW cells expressing CFP-Rab5 or CFP-Rab5(Q79L), as indicated. For each condition, 3 independent experiments were quantified, with ≥ 10 cells per replicate. Data are means ± SEM. **g.** Comparison of Rab7 FRAP in LRRK2^+/+^ (left) versus LRRK2^−/−–^ RAW cells (right). FRAP was measured as described in untreated or ΔpH-dissipated LRRK2^+/+^ or LRRK2^−/−^ RAW cells expressing GFP-Rab7 or GFP-Rab7(Q67L), and the vesicle Rab7 fluorescence recovery monitored. For each condition, 3 independent experiments were quantified, with ≥ 10 cells per replicate. Data are means ± SEM.

### LRRK2 is required for the pH-dependent insensitivity of Rab GTPases to GAP function

The availability of LRRK2-deficient cells enabled us to test whether the kinase is indeed responsible for the inhibition of GAP activity recorded in cells where the pH gradients had been dissipated. As discussed above, the rate and extent of recovery following photobleaching of GFP-tagged Rab proteins can be used to estimate their rate of dissociation from organelles, which is indicative of the rate of GTP hydrolysis. Therefore, we compared the rate of FRAP of Rab5 and Rab7 between LRRK2^+/+^ and LRRK2^−/−^ macrophages. While dissipating the ΔpH depressed the rate and extent of FRAP in the LRRK2^+/+^ cells (left panel in Fig. 6f, as reported also in Fig. 3a), no change was detectable in the LRRK2^−/−^ cells (right panel in Fig. 6f). Importantly, as in wildtype (LRRK2^+/+^) macrophages (left panel in Fig. 6f, as reported also in Fig. 3a), the recovery of constitutively-active (Q79L) Rab5 was markedly slower than that of wildtype Rab5 in the LRRK2-knockout cells (right panel in Fig. 6f), validating the FRAP analysis approach in the LRRK2^−/−^ cells. Essentially identical results were obtained when LRRK2^+/+^ and LRRK2^−/−^ macrophages were transfected with wildtype and constitutively active (Q67L) Rab7 (left and right panels of Fig. 6g, respectively). Based on these data we concluded that the pH-induced Rab activation requires LRRK2. When this kinase is deleted, the GTPases appear to be insensitive to the luminal pH and their cognate GAPs are able to function normally to promote Rab-GTP hydrolysis.

## DISCUSSION

Dissipation of the endomembrane pH gradient has long been reported to arrest vesicular traffic ^8^. Multiple stages of the endocytic and secretory pathways are altered when the luminal pH is neutralized and, as a consequence, a variety of signaling and scaffolding molecules have been implicated. In the secretory pathway COPI-dependent membrane budding was found to require luminal acidification ^18–21^ and, because of its role in COPI assembly, ARF1 was suspected to contribute to the pH sensitivity. Analogously, ARF6 and its exchange factor ARNO that are required for some forms of endocytosis and vesicle uncoating were shown to respond to dissipation of the luminal acidification ^52^. Interestingly, these bind to the V-ATPase in a pH-dependent manner ^53^ Most recently, luminal acidification was found to signal the early-to-late endosome transition by prompting the dissociation of the Vps34 complex from membranes and the consequent cessation of PtdIns(3)P synthesis ^54^. Whether these are all independent and unrelated effects remains unknown; no unifying mechanism has been identified.

Here, we describe experiments indicating that the luminal pH of endosomes dictates the state of activation of endocytic Rab GTPases, including Rab5 and Rab7. Upon dissipation of existing transmembrane pH gradients, these GTPases became abnormally activated. We demonstrated that in cells where the ΔpH had been dissipated Rab5 and Rab7 were hyperactive and remained GTP-locked on enlarged endocytic compartments. Likely these enlarged compartments are the result of stalled endocytic maturation, as suggested by the inability of internalized dextran to reach the late compartments of the pathway (see Fig. 1f-g), resulting from ongoing fusion in the absence of the normal vesiculation and tubulation mechanisms needed to resolve endosomal contents. Indeed, Rab5 and Rab7 cycling is required for these mechanisms ^55,56^.

Stimulation of Rab5 and Rab7 was accompanied by their slower dissociation from endosomal membranes, indicative of reduced rates of GTP hydrolysis. Because Rab GTPases have poor intrinsic GTPase activity, this suggested that the recruitment or activity of their cognate GAPs was affected. In the case of Rab5, we found that two putative GAP proteins – RabGAP-5 ^57^ and TBC1D2B ^58^– remained associated with endosomes following pH dissipation. These data implied that the ability of the GAPs to induce hydrolysis, rather than their recruitment, was affected by the ΔpH. Moreover, the observation that multiple Rabs were impacted, suggested the existence of a more general, common mechanism whereby pH regulates their ability to hydrolyse GTP.

Recent studies have revealed that phosphorylation of the switch-II region of Rab proteins blocks their ability to interact with regulatory proteins such as GDIs and GAPs ^37–39,59^. The phospho-Rab proteins are consequently trapped in the GTP-bound state, remaining attached and hyperactive on endomembranes ^37,60^. Such Rab phosphorylation occurred in cells where pH gradients were dissipated and was shown by pharmacological and genetic means to be exerted by LRRK2, a kinase associated with dominantly-inherited and sporadic Parkinson’s disease ^61–64^. Under these conditions the kinase itself underwent phosphorylation at a site (S1292) that is indicative of its activation. The exact mechanism of LRRK2 stimulation has been unclear; diverse reports have attributed activation to conceptually disparate signals including lysosomal “stress”, endomembrane damage and cell exposure to lysosomotropic drugs or ionophores ^45–48^. A unifying feature of all these agents and conditions is the neutralization of luminal pH. We therefore propose that dissipation of the transmembrane pH gradient, which was similarly and purposefully induced in our experiments by treatment with concanamycin/nigericin, is the immediate cause of LRRK2 recruitment and activation on endomembranes.

In addition to becoming catalytically more active, LRRK2 was recruited to endomembranes in response to concanamycin/nigericin. It remains unclear how LRRK2, a cytoplasmic enzyme, senses the dissipation of the luminal acidification. We envisage the existence of a transmembrane transducer that conveys information of the luminal pH to LRRK2 across the bilayer. Interestingly, LRRK2 has been shown to interact with the *a* subunit of the V-ATPase ^65^, which spans the bilayer and has been hypothesized to be a transmembrane pH sensor ^66,67^. Moreover, several regulators of endocytosis, including ARNO and Arf6, have been shown to bind to the *a* subunit in a pH-dependent manner ^52,53,68–70^, reinforcing the possibility that LRRK2 may bind to the V-ATPase similarly. However, other possibilities should be considered: LRRK2 activation has been tied to the recruitment to membranes of Rab29 in some studies ^71,72^. Whether recruitment of this Rab is the cause or a consequence of LRRK2 activation is unclear. In our system, however, Rab29 was not visibly recruited to sites of LRRK2 accumulation on endomembranes upon ΔpH dissipation (data not shown).

Activating mutations in LRRK2 are one of the most common genetic causes of familial and sporadic Parkinson’s disease ^73,74^. In Parkinson’s these mutations result in misregulation of intracellular traffic and lysosomal degradation pathways ^37,75–80^. Vast changes in vesicle positioning and endocytic continuity are observed when LRRK2 is hyperactive ^81^, although the precise mechanism responsible for these changes is incompletely understood. Because LRRK2 phosphorylates a subgroup of Rab proteins ^37,59,78^, altered functionality of these GTPases may underlie the endocytic alterations. Our data, which revealed that LRRK2 confers Rab5 and Rab7 refractoriness to their corresponding GAPs, may contribute to understanding the etiology of Parkinson’s disease. It is noteworthy that, while we demonstrated that Rab8 and Rab10 are directly phosphorylated by LRRK2 when luminal pH is altered, technical limitations have prevented us from establishing unequivocally that Rab5 and 7 are similarly phosphorylated. Nevertheless, it is clear that LRRK2-mediated phosphorylation is responsible for their reduced GTPase activity, deduced from their enhanced effector binding and inability to dissociate from early and late endosomes, respectively. An indirect effect, attributable to phosphorylation of the GAPs themselves or of ancillary proteins cannot be discounted.

In sum, our data suggest that LRRK2 responds to changes in luminal pH, increasing its ability to phosphorylate Rab GTPases and possibly other proteins involved in endocytic traffic. We propose that by dictating the state of activation of LRRK2 and in turn that of Rab GTPases, the development of a progressive luminal acidification serves as a timing device to control maturation of the endocytic pathway. Transitions between maturation stages may only be enabled when LRRK2 is sufficiently inhibited by the developing acidification to permit the termination of the activity of the Rab responsible for the preceding stage. This would position LRRK2 as a key regulator of endocytic recycling and maturation.

## Data availability

Data supporting the findings of this study are available from the corresponding author on reasonable request. Source data are provided with this paper.

## Acknowledgements

We thank Fabienne Geiger and Sahel Taei for their superb experimental assistance. This work was supported by Canadian Institutes of Health Research (FDN-143202). MM was the recipient of a Heart and Stroke Foundation of Canada (HSFC) Pfizer Research Fellowship and a Roifman Scholar Award for Young Investigators.

## Author contributions

Conceptualization: MM, SG; Methodology: MM, KH, SG; Formal analysis: MM, KH; Investigation: MM, KH; Resources: SG; Writing - original draft: MM, SG. Writing - review & editing: MM, KH, SG; Supervision: SG

## MATERIALS AND METHODS

### Cell culture

RAW 264.7, RAW 264.7 *LRRK2^−/−^* and HeLa cells were obtained from and authenticated by the American Type Culture Collection (ATCC). AD293 cells were obtained from and authenticated by Stratagene. All cell lines tested negative for mycoplasma contamination by DAPI staining. RAW cell lines were grown in RPMI-1640 medium containing L-glutamine and 10% heat-inactivated fetal calf serum (FCS; MultiCell, Wisent), at 37°C under 5% CO_2_. HeLa and AD293 cells were grown in DMEM containing L-glutamine and 10% heat-inactivated FCS at 37°C under 5% CO_2_.

### Reagents

Mammalian expression vectors were obtained from the following sources: CFP-Rab5A ^82^; GFP-Rab5A ^83^; RFP-Rab5A ^84^; CFP-Rab5A(Q79L) ^82^; GFP-Rab5A(Q79L) (35140, Addgene); mCherry-Rab5A(Q79L) (35138, Addgene); CFP-Rab5A(S34N) ^82^; RFP-Rab7A ^85^; RFP-Rab7A(Q67L) ^85^, RFP-Rab7A(T22N) ^85^; GFP-Rab7A ^86^; GFP-Rab7A(Q67L) ^86^; GFP-Rab10A (49472, Addgene); GFP-Rab8A (24898, Addgene), R5BD-YFP ^87^, GFP-APPL1 ^88^; GFP-RILP_C33_ ^36^; RabGAP5-GFP ^57^; TBC1D2B-GFP (provided by Dr. Christian Rocheleau, McGill University); LRRK2-neonGreen (provided by Drs. Luis Bonet-Ponce and Mark Cookson, National Institute of Health).

Primary antibodies were purchased from the following vendors: LRRK2 (ab133474, Abcam) LRRK2 pS1292 (ab203181, Abcam); Rab10 (ab230260, Abcam), Rab10 pT73 (ab241060, Abcam); Rab8 (ab241061, Abcam), Rab8 pT72 (ab230260, Abcam), vinculin (MAB3574, Sigma-Aldrich); α-tubulin (T5168, Sigma-Aldrich). Horseradish peroxidase-conjugated secondary antibodies were purchased from Jackson ImmunoResearch Labs.

Concanamycin A, nigericin, sulforhodamine B and chloroquine were purchased from Sigma-Aldrich. FM4-64 and leucyl-L-leucine methyl ester (LLOMe) were from Thermo Fisher Scientific. MLi-2 was purchased from Abcam. Fluorescent conjugated 10 kDa dextrans were purchased from Invitrogen. Paraformaldehyde (PFA; 16% wt/vol) was from Electron Microscopy Sciences.

### Transient DNA transfection

RAW cell lines were plated on 18 mm glass coverslips at ∼2×10^5^ cells·mL^−1^, 16-24 h prior to transfection. FuGENE HD (Promega) transfection reagent was used to transfect RAW cells at a 3.5:1 ratio (using 1.75 µL FuGENE HD and 0.5 µg DNA per well). HeLa or AD293 cells were plated at on 18 mm glass coverslips at a concentration of ∼5×10^4^ or ∼1×10^5^ cells·mL^−1^, respectively, 16-24 h prior to transfection. FuGENE 6 (Promega) transfection reagent was used according to the manufacturer’s instructions to transfect at a 3:1 ratio (using 1.5 µL FuGENE 6 and 0.5 µg DNA per well). In all cases, monolayers were used for experiments 16 h after transfection.

### pH dissipation experiments

Cells were placed in isotonic K^+^ medium (15 mM HEPES, 140 mM KCl, 5 mM glucose, 1 mM MgCl_2_, and 1 mM CaCl_2_) with or without 250 nM concanamycin A and 5 µg·mL^−1^ nigericin, and incubated for 30 min or 60 min at 37°C. In some experiments, endocytic compartments were labeled with fluorescently conjugated dextran, FM4-64 or sulforhodamine B. To assess fusion of early endosomes with late endosomes/lysosomes, cells were first pulsed with 100 µg·mL^−1^ Alexa 488-dextran for 16 h followed by 1 h chase to label exclusively late endosomes/lysosomes. To label early endosomes cells were then pulsed with 0.5 mg·mL^−1^ tetramethylrhodamine-dextran for 30 min and chased for the indicated times under control or pH-dissipating conditions. After incubations, cells were washed 3× with 1× PBS, and viewed live. For FM4-64 labeling, the cells were first incubated in 1 µM FM4-64 for 10 or 20 min to label early or late endocytic compartments, respectively. After incubations, cells were washed 3× with 1× PBS, used for pH dissipation experiments, and viewed live. For sulforhodamine B labeling, cells were either incubated with 30 µg·mL^−1^ sulforhodamine B during 30 min pH dissipation, or incubated for 20 min with 30 µg·mL^−1^ sulforhodamine B before 30 min pH dissipation, to label early or late endocytic compartments, respectively. After pH dissipation, cells were washed 3× with 1× PBS cells and viewed live. To validate its effectiveness as an endolysosome-disrupting agent, sulforhodamine B-loaded cells were treated with 400 µM LLOMe for 30 min. In some cases, monolayers were first pretreated with 50 nM MLi-2 for 90 min at 37°C in isotonic K^+^ medium prior to pH dissipation experiments. Unless viewed live, following pH dissipation, monolayers were fixed in 3% paraformaldehyde for 10 min at room temperature, then washed with PBS (MultiCell).

### Immunoblotting

The day before experiments, RAW cells were seeded in T-25 flasks at a concentration of ∼4×10^6^ per flask. After pH dissipation experiments, monolayers were lysed using RIPA buffer (Sigma-Aldrich) containing protease and phosphatase inhibitors (Pierce). Protein concentrations were determined using BCA protein assays (Pierce). 20-40 µg of protein was then diluted in Laemmli buffer (BioRad) and loaded per well on 4-15% gradient polyacrylamide gels (BioRad) for SDS-PAGE analysis. The gels were then transferred to polyvinylidene difluoride or nitrocellulose membranes and treated in blocking buffer consisting of TBS with 5% BSA and 0.05% Tween-20 for 1 h at room temperature. Primary antibody staining was in blocking buffer for 1 h at room temperature. Primary antibody dilutions were as follows: LRRK2 (1:1000), LRRK2 pS1292 (1:1000), Rab10 (1:1000), Rab10 pT73 (1:1000); Rab8 (1:1000), Rab8 pT72 (1:1000), vinculin (1:2000); α-tubulin (1:5000). After washing the membranes in TBS containing 0.05% Tween-20, samples were incubated 1 h at room temperature with appropriate HRP-conjugated secondary antibodies at 1:3000 dilution. Blots were developed using the ECL Prime Western Blot detection reagent (Cytiva Amersham) and visualized with a Bio-Rad ChemiDoc MP Imaging System and Image Lab software 5.2.1.

### Microscopy and image analyses

Confocal images were acquired using a spinning disk system (Quorum Technologies Inc.). The instrument consists of a microscope (Axiovert 200M; Zeiss), scanning unit (CSU10; Yokogawa Electric Corporation), electron-multiplied charge-coupled device camera (C9100-13; Hamamatsu Photonics) or ORCA-Fusion BT Digital CMOS camera (C15440-20UP; Hamamatsu Photonics), five-line (405-, 443-, 491-, 561-, and 655-nm) laser module (Spectral Applied Research), and filter wheel (MAC5000; Ludl) and is operated by Volocity v6.3. Images were acquired using a 63× oil objective (Zeiss), with an additional 1.5× magnifying lens and the appropriate emission filter. For live experiments, cells were maintained at 37°C using an environmental chamber (Live Cell Instruments). Image processing and background-subtracted fluorescence analyses, including the calculation of Pearson’s colocalization coefficients (r), were performed using Volocity v6.3. Image deconvolution was done on acquired Z-stacks within the Volocity Restoration module, using the iterative restoration function. Calculated fluorochrome point spread functions were used to deconvolve individual channels for 5-8 iterations, until a confidence limit of >90% was achieved.

Confocal Z stacks were used for Statistical Object Distance Analysis (SODA) in the open-source analysis software Icy (http://icy.bioimageanalysis.org/), to quantitatively characterize the spatial positioning of Rab5A/Rab7A puncta with LRRK2 puncta within the cell. ∼750 nm deep image stacks of cells expressing transiently expressing fluorescent Rab5A or Rab7A and LRRK2 were analyzed using the Icy SODA 2 color protocol ^44^. SODA used the Ripley’s K function and statistical thresholding relative to a random distribution (null hypothesis) to quantify the coupling probability of spots identified between two emission channels, using sequential concentric rings centered around the center of intensity of channel 1 spots (in this case, Rab5A/Rab7A endosomes). The protocol included spot detection, using the Wavelet Spot Detector block (Scale 2), and cell ROI selection by binary cell mask, using the Thresholder block. SODA analysis was limited to a 550 nm radius (5 pixels) from the identified Rab5A/Rab7A endosomes. Statistics for single and coupled spots within individual cells, mean coupling probabilities and coupling *p-value*s were extracted from the analysis.

Fluorescence Lifetime Imaging Microscopy (FLIM) was performed on live control or ΔpH-dissipated cells at 37°C using an Olympus IX81 microscope equipped with a Lambert FLIM module and a Hamamatsu EM-CCD. Cells previously transfected with CFP-Rab5A/Rab5A(Q79L) (donor fluorophore), with or without YFP-R5BD (acceptor fluorophore) were imaged with a 60× oil objective. For FLIM quantifications, CFP fluorescence lifetimes (τ) in the presence (donor + acceptor) and absence (donor only) of YFP acceptor were measured. Images were exported and analyzed using LI-FLIM v1.2.10.35 software. Förster resonance energy transfer (FRET) efficiencies (E) were calculated from the formula:

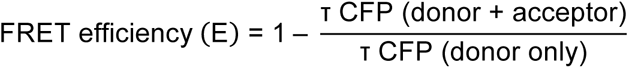

Fluorescence recovery after photobleaching (FRAP) experiments were conducted on live control or ΔpH-dissipated cells at 37°C using an A1R point-scanning confocal system (Nikon). Images were acquired using a 63× oil objective, 1.2-AU pinhole, Galvano scanning mode with no line averaging, at 256 pixel resolution and with 1.83× rectangular zoom. For a single field, after 5 s of initial imaging, a region of interest ∼2 µm in diameter was bleached for 1 s using the 405 laser at 100% power, followed by imaging for 2 min for fluorescence recovery. Images were exported and analyzed for fluorescence intensity using Volocity v6.3 software. After background subtraction, fluorescence intensity units were normalized to a non-bleached vesicle and transformed to a 0-1 scale to correct for differences in bleaching depth and to allow for comparison of up to 30 individual FRAP curves per condition ^89^. Normalizations were done using Microsoft Excel software. Graphpad Prism software was used to fit the FRAP curves to a single exponential, plotted as fractional recovery over time. Values derived from this non-linear regression were used to calculate half-time of recovery (t_1/2_) and mobile fraction ^90^.

Differential interference contrast microscopy (DIC) was performed on control or ΔpH-dissipated cells using Leica DM IRE2 inverted fluorescence microscope with a 100× oil objective and digital images were acquired with an ORCA-ER camera (Hamamatsu) controlled by Open Lab V3.4 (Improvision) software.

For transmission electron microscopy (TEM) analyses, control or ΔpH-dissipated cells were fixed at room temperature in 2% glutaraldehyde in 0.1 M phosphate buffer (pH 7.0) for 1 h. Samples were then stained *en bloc* for 1h with 4% uranyl acetate, dehydrated through increasing alcohol concentrations and embedded in Epon. 40-50 nm sections were cut on a Leica Ultracut R, counterstained with 2% lead citrate and finally observed at 60-80 kV in a Philips CM10 transmission electron microscope.

### General methodology and statistics

Data calculations and normalizations were done using Microsoft Excel 2011 (Microsoft Corporation) or GraphPad Prism v9.4 software (GraphPad Software, Inc.). Because experiments were, for the most part, *in vitro* imaging determinations of individual cells, samples were assigned to groups according to specific experimental treatments (control vs. experimental group). The number of individual experiments and the number of determinations per experiment were selected to attain an estimate of the variance compatible with the statistical tests used, primarily student’s *t* test or one-way ANOVA. Each type of experiment was performed a minimum of three separate times (biological replicates) and a minimum of ten individual event determinations (equivalent, but not identical to technical replicates). Data was tested for normality using the D’Agostino-Pearson test, and appropriate testing applied. No data were excluded as outliers. All statistics were calculated using GraphPad Prism v9.4.

## EXTENDED DATA FIGURE LEGENDS

**Extended Data Figure 1.**
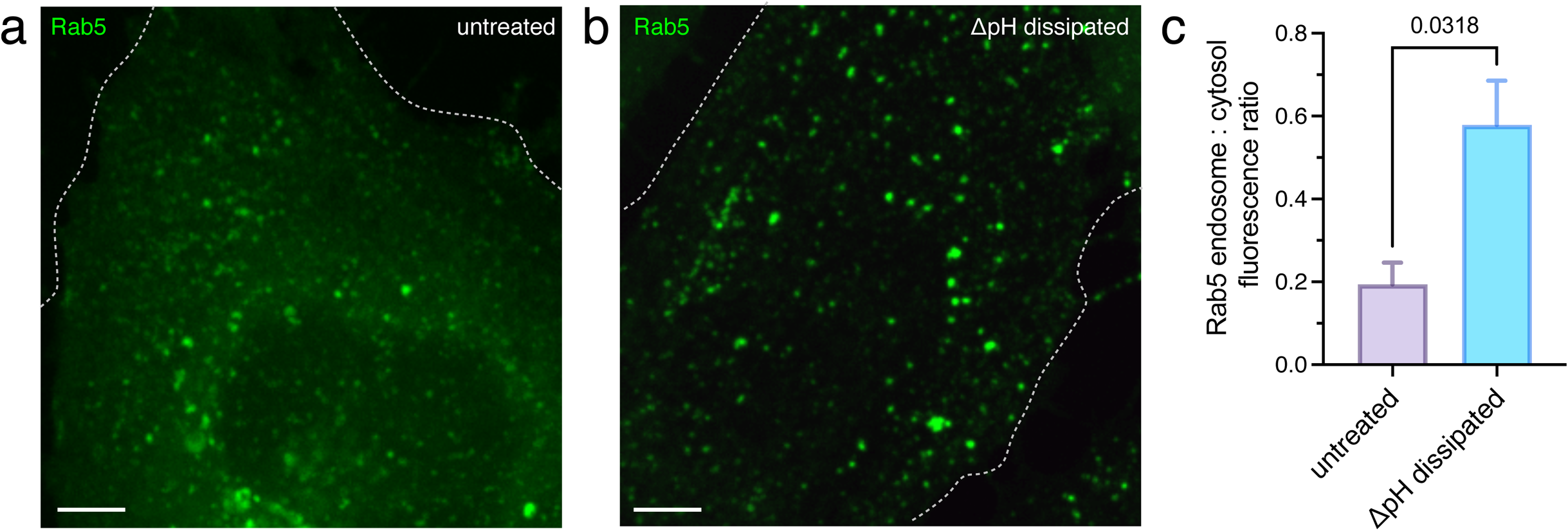
ΔpH dissipation induces endosome enlargement in HeLa cells. **a**-**b** Untreated (**a**) and ΔpH-dissipated (**b**) HeLa cells expressing GFP-Rab5 (green). **c.** Quantitation of the ratio of membrane-associated (punctate) to cytoplasmic (diffuse) signal of GFP-Rab5 in transiently transfected untreated and ΔpH-dissipated HeLa cells. For each condition, 3 independent experiments were quantified, with ≥ 10 cells per replicate. Data are means ± SEM. All images are representative of ≥ 30 fields from ≥ 3 experiments of each type. Scale bars: 5 µm.

**Extended Data Figure 2.**
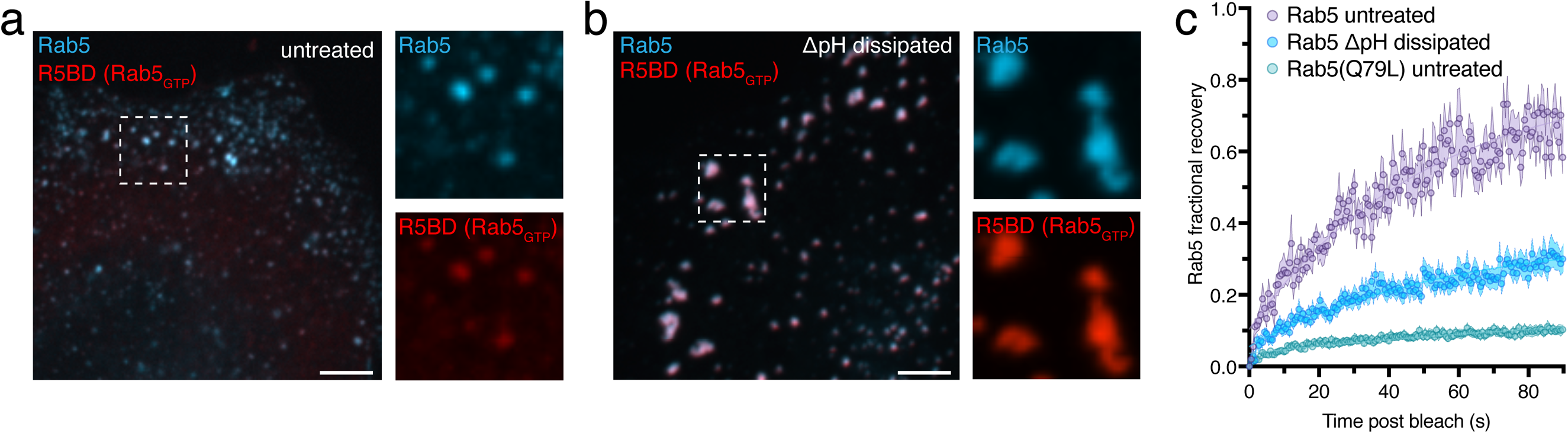
Hyperactivation of Rab GTPases during ΔpH dissipation occurs also in HeLa cells. **a.** Untreated HeLa cells expressing Rab5 (cyan) and R5BD (red), a biosensor for GTP-bound Rab5. Side panels in **a** and **b** show individual Rab5 and R5BD channels at 2.5× magnification. **b.** ΔpH-dissipated HeLa cells expressing Rab5 (cyan) and R5BD (red). Images in **a** and **b** are representative of ≥ 30 fields from ≥ 3 experiments of each type. Scale bars: 5 µm. **c.** The fluorescence of individual vesicles expressing GFP-Rab5 or GFP-Rab5(Q79L) was photobleached in untreated or ΔpH-dissipated HeLa cells and the rate and extent of fluorescence recovery was monitored. For FRAP curves, average vesicle fluorescence before photobleaching was defined as 1.0, and fluorescence recovery normalized to that of an unbleached vesicle to correct for bleaching incurred during the acquisition of images during the recovery phase. For each condition, 3 independent experiments were quantified, with ≥ 10 cells per replicate. Data are means ± SEM.

**Extended Data Figure 3.**
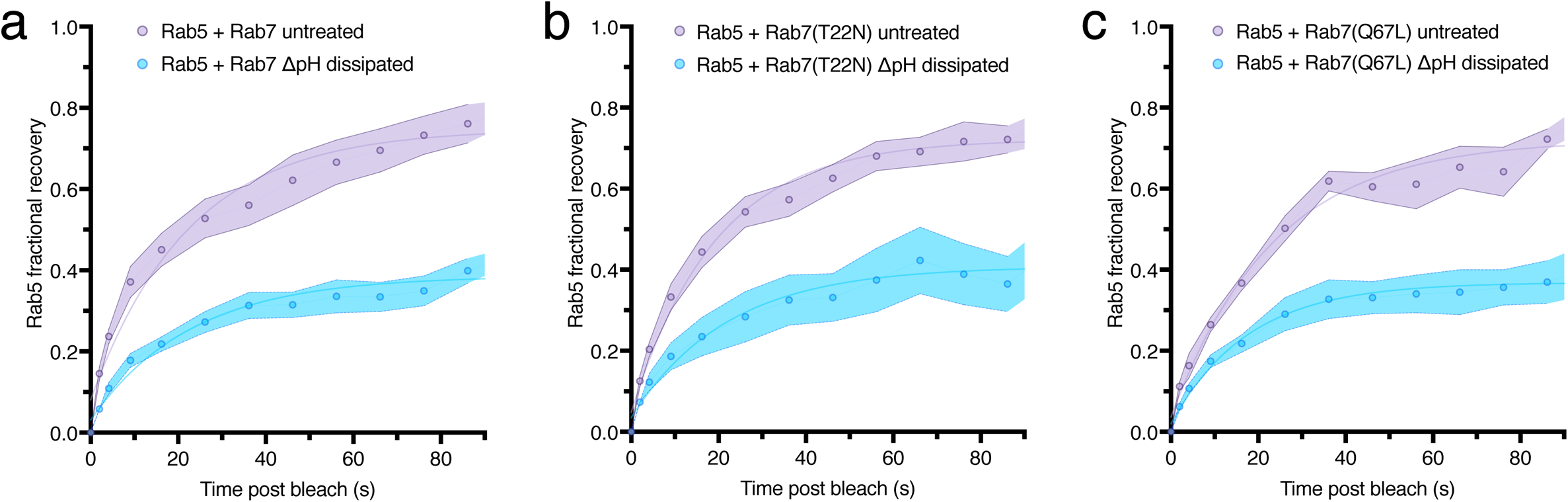
Activation of Rab GTPases during ΔpH dissipation is not the result of impaired traffic between the early and late endocytic compartment. Wildtype CFP-Rab5 was co-expressed in RAW cells co- with RFP-Rab7 (**a**), RFP-Rab7(T22N) (**b**) or RFP-Rab7(Q67L) (**c**). Individual CFP-Rab5-positive vesicles were photobleached in untreated or ΔpH-dissipated cells and fluorescence recovery monitored over 90 s. For each condition, 3 independent experiments were quantified, with ≥ 10 cells per replicate. Data are means ± SEM.

**Extended Data Figure 4.**
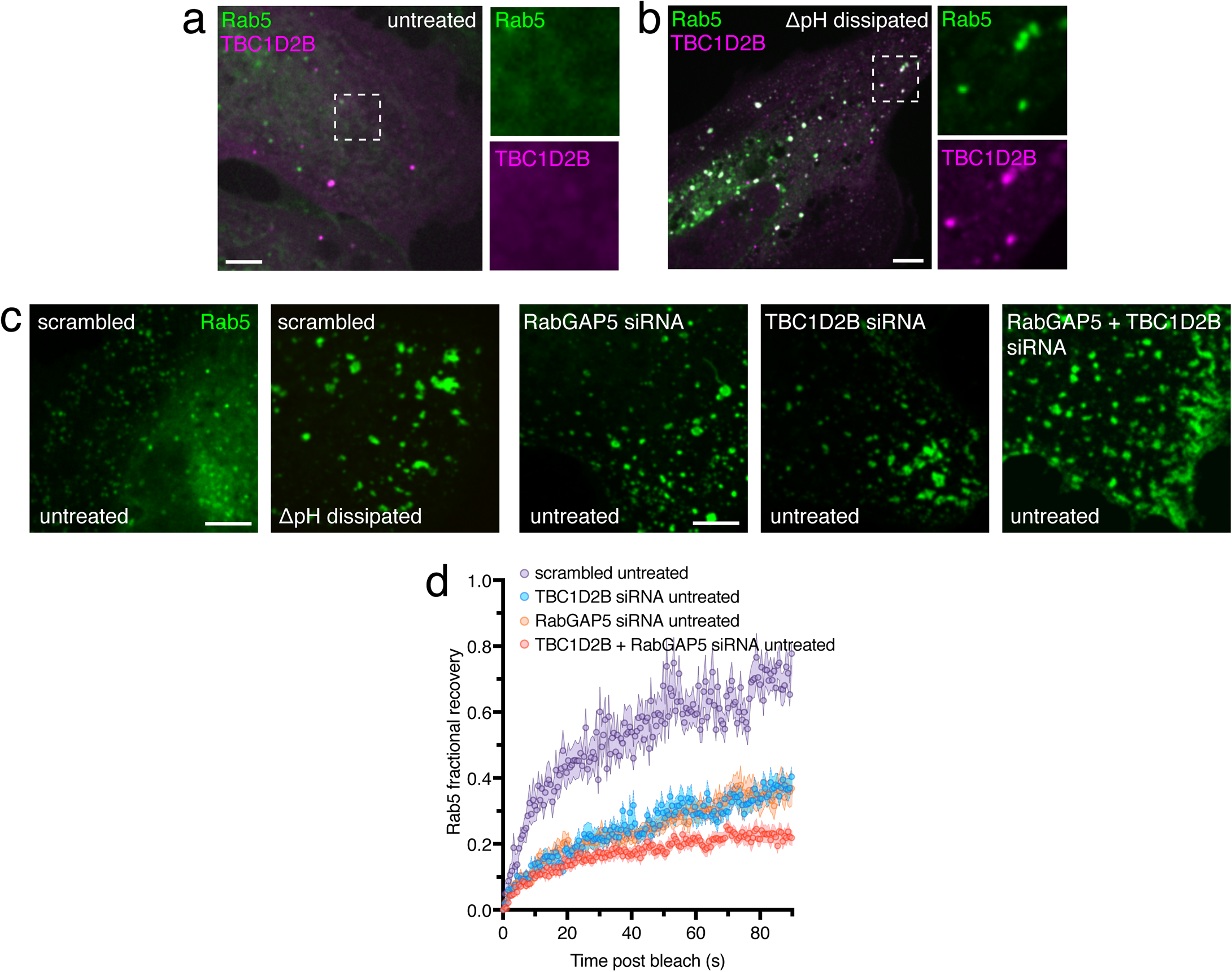
Rab5-GAPs inactivate Rab5 on endosomes. **a.** Untreated HeLa cells expressing Rab5 (green) and TBC1D2B (magenta). Side panels show individual Rab5 and TBC1D2B channels at 2.8× magnification. **b.** ΔpH-dissipated HeLa cells expressing Rab5 (green) and TBC1D2B (magenta). Side panels show individual Rab5 and TBC1D2B channels at 2.9× magnification. **c.** Visualization of Rab5-positive compartments (green) in untreated or ΔpH-dissipated HeLa cells that had been previously electroporated in the presence of scrambled, RabGAP5 or TBC1D2B siRNA, as indicated. All images are representative of ≥ 30 fields from ≥ 3 experiments of each type. All scale bars: 5 µm. **d.** Individual Rab5-positive vesicles were photobleached in HeLa cells expressing GFP-Rab5 (green) that had been previously electroporated in the presence of scrambled, RabGAP5, TBC1D2B or RabGAP5+TBC1D2B siRNA, as indicated. GFP-Rab5 fluorescence recovery was then monitored. For each condition, 3 independent experiments were quantified, with ≥ 10 cells per replicate. Data are means ± SEM.

**Extended Data Figure 5.**
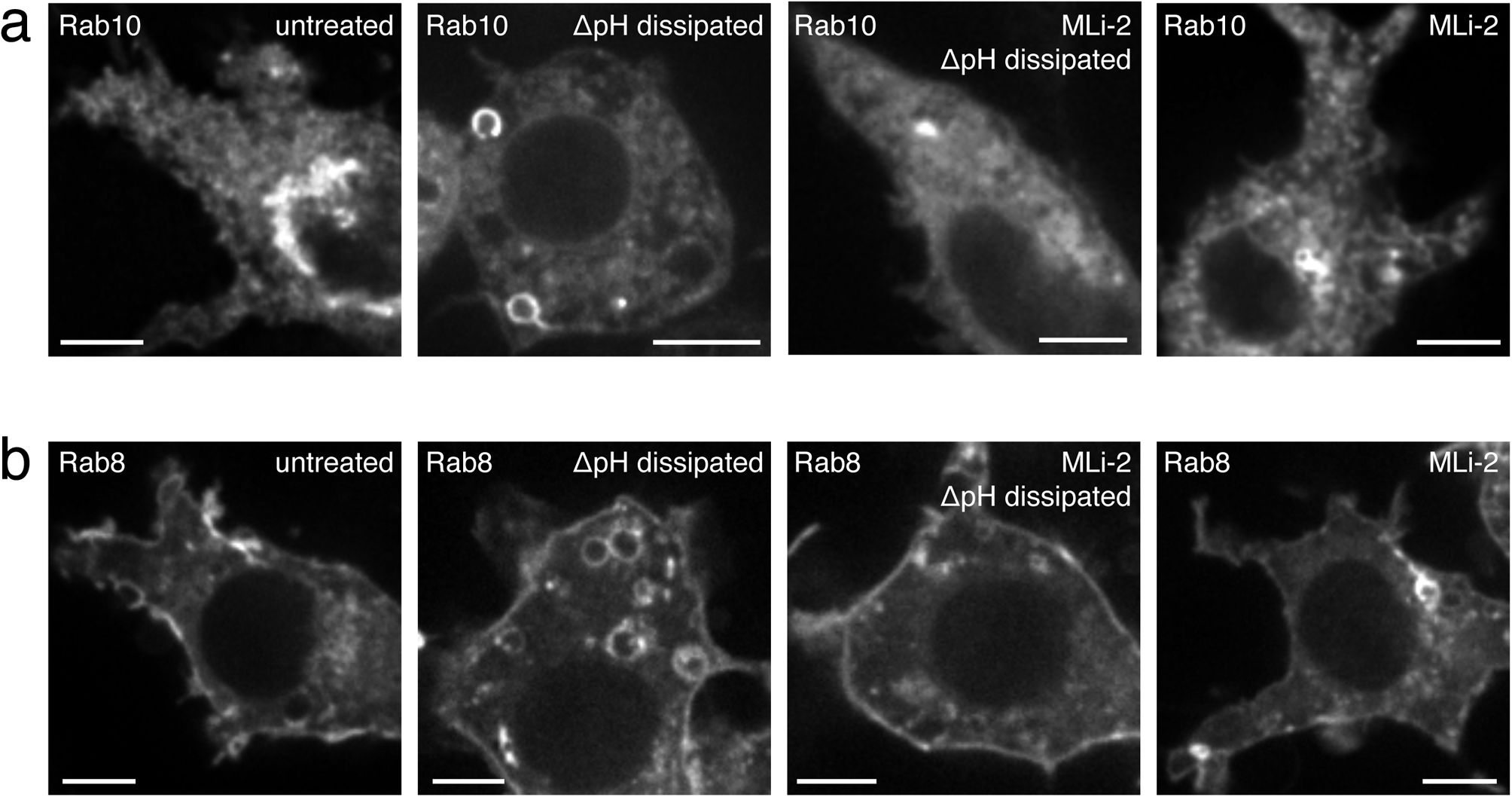
LRRK2 activity is required for the enlargement of Rab10-positive and Rab8-positive vesicles induced by ΔpH dissipation. **a.** Effect of ΔpH dissipation and of MLi-2 on the appearance of Rab10-positive compartments (white) of RAW cells. **b.** Effect of ΔpH dissipation and of MLi-2 on the appearance of Rab8-positive compartments (white). All images are representative of ≥ 30 fields from ≥ 3 experiments of each type. Scale bars: 5 µm.

**Extended Data Figure 6.**
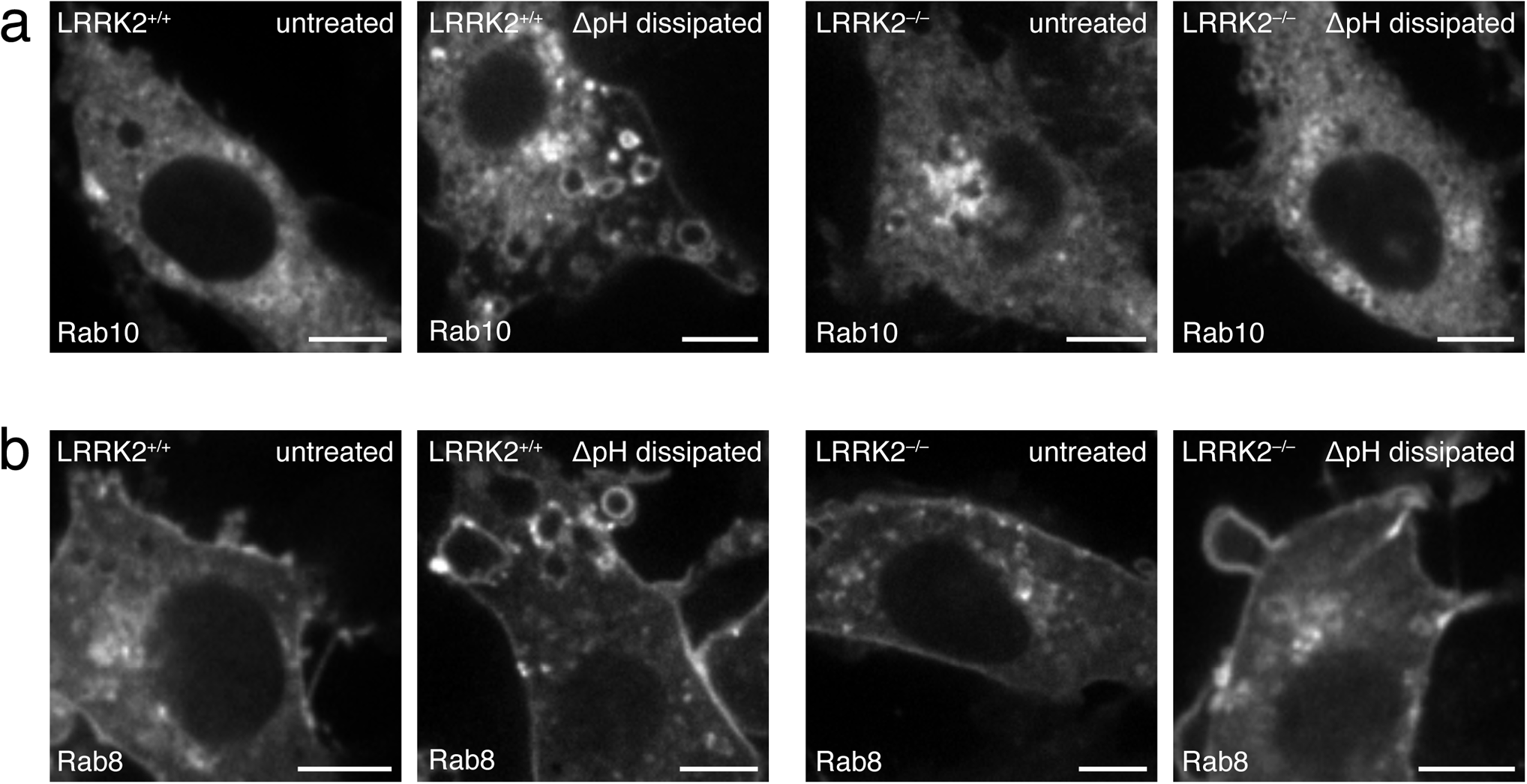
LRRK2^−/−^ RAW cells do not form enlarged Rab10- or Rab8-positive vacuoles after ΔpH dissipation. **a.** Effect of knocking out LRRK2 on the vacuolation of Rab10-positive membranes (white) induced by the ΔpH dissipation in RAW cells. **b.** Effect of knocking out LRRK2 on the vacuolation of Rab8-positive membranes (white) induced by the ΔpH dissipation. All images are representative of ≥ 30 fields from ≥ 3 experiments of each type. Scale bars: 5 µm.

**Source Data Figure 4:**
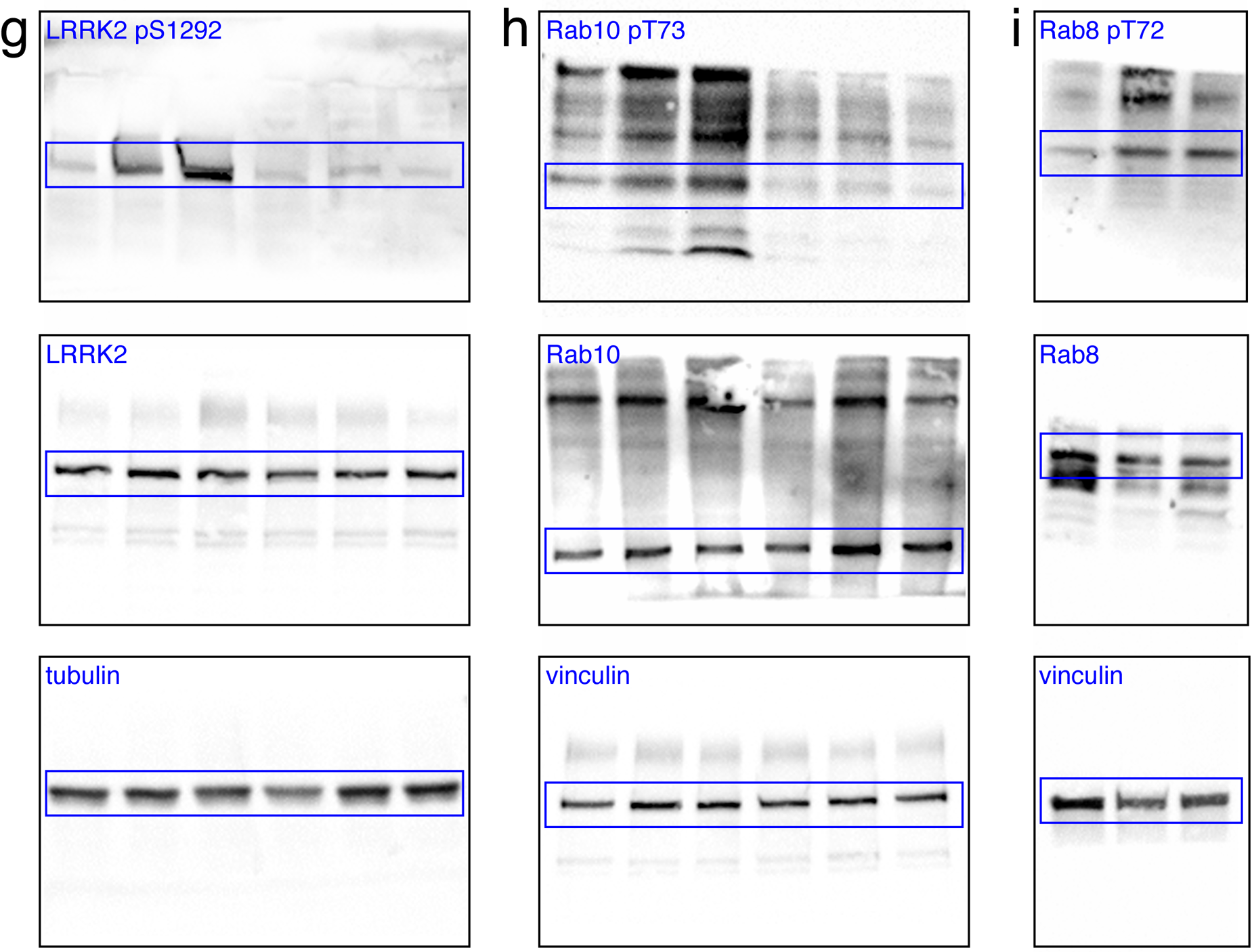
Full size immunoblots corresponding to Fig. 4 **g**, **h**, **i**. Bands shown in main figures are highlighted by blue boxes.

**Source Data Figure 6:**
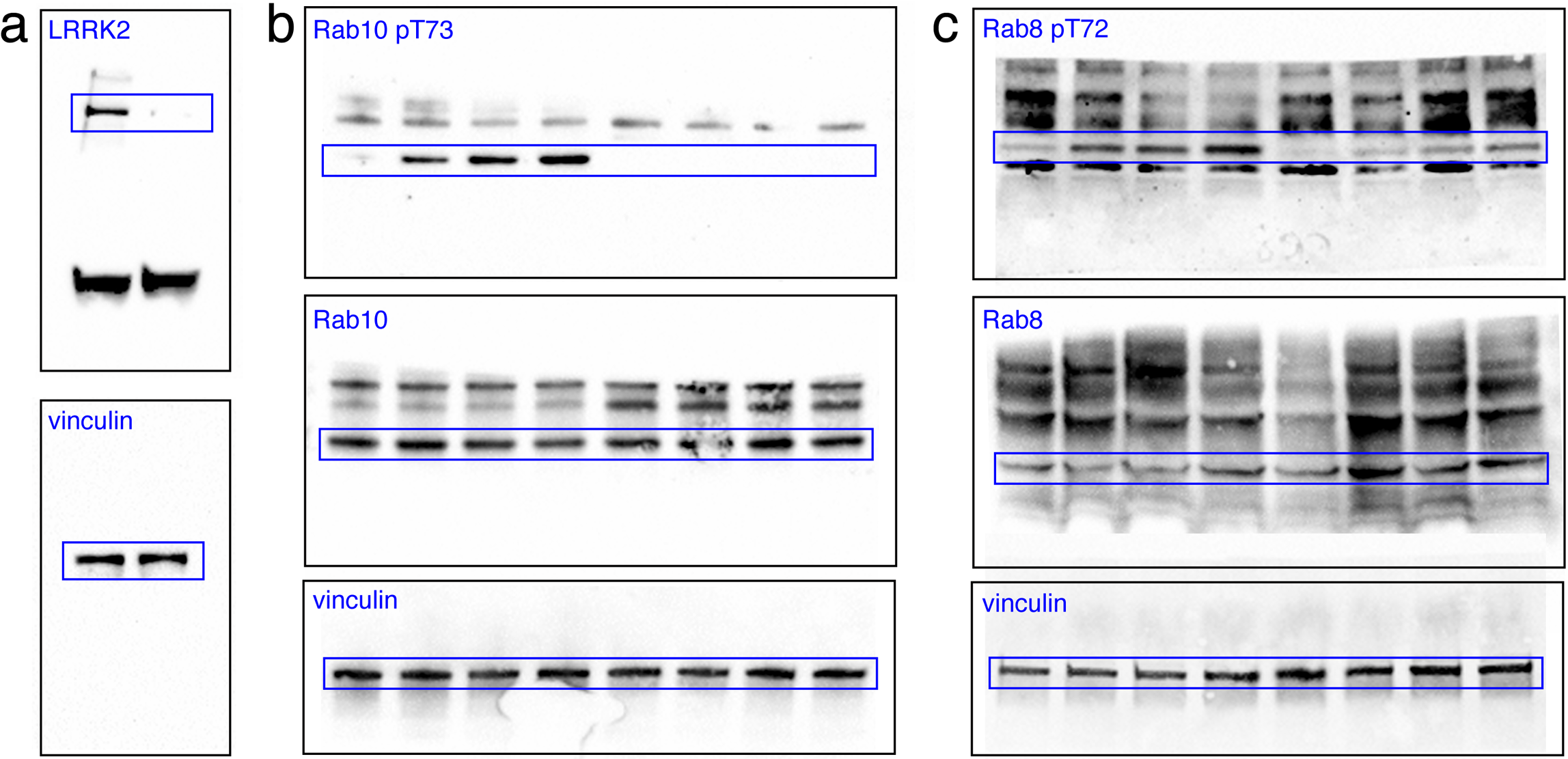
Full size immunoblots corresponding to Fig. 6 **a**, **b**, **c**. Bands shown in main figures are highlighted by blue boxes.

## REFERENCES

1. Maxfield, F. R. & Yamashiro, D. J. Endosome acidification and the pathways of receptor-mediated endocytosis. Adv Exp Med Biol 225, 189–198 (1987).

2. Tycko, B. & Maxfield, F. R. Rapid acidification of endocytic vesicles containing alpha 2-macroglobulin. Cell 28, 643–651 (1982).

3. Chadwick, S. R., Grinstein, S. & Freeman, S. A. From the inside out: Ion fluxes at the centre of endocytic traffic. Current Opinion in Cell Biology 71, 77–86 (2021).

4. Yamashiro, D. J., Fluss, S. R. & Maxfield, F. R. Acidification of endocytic vesicles by an ATP-dependent proton pump. J Cell Biol 97, 929–934 (1983).

5. Schneider, D. L. Membranous localization and properties of ATPase of rat liver lysosomes. The Journal of membrane biology 34, 247–261 (1977).

6. Schneider, D. L. A membranous ATPase unique to lysosomes. Biochemical and Biophysical Research Communications 61, 882–888 (1974).

7. Collins, M. P. & Forgac, M. Regulation and function of V-ATPases in physiology and disease. Biochim Biophys Acta Biomembr 1862, 183341 (2020).

8. Weisz, O. A. Acidification and Protein Traffic. in International Review of Cytology vol. 226 259–319 (Elsevier, 2003).

9. Davis, C. G. et al. Acid-dependent ligand dissociation and recycling of LDL receptor mediated by growth factor homology region. Nature 326, 760–765 (1987).

10. Yamashiro, D. J., Tycko, B., Fluss, S. R. & Maxfield, F. R. Segregation of transferrin to a mildly acidic (pH 6.5) para-Golgi compartment in the recycling pathway. Cell 37, 789–800 (1984).

11. Dautry-Varsat, A., Ciechanover, A. & Lodish, H. F. pH and the recycling of transferrin during receptor-mediated endocytosis. Proc Natl Acad Sci U S A 80, 2258–2262 (1983).

12. Schwartz, A. L., Bolognesi, A. & Fridovich, S. E. Recycling of the asialoglycoprotein receptor and the effect of lysosomotropic amines in hepatoma cells. J Cell Biol 98, 732–738 (1984).

13. De Duve, C., Pressman, B. C., Gianetto, R., Wattiaux, R. & Appelmans, F. Tissue fractionation studies. 6. Intracellular distribution patterns of enzymes in rat-liver tissue. Biochem J 60, 604–617 (1955).

14. Essner, E. & Novikoff, A. B. Localization of acid phosphatase activity in hepatic lysosomes by means of electron microscopy. J Biophys Biochem Cytol 9, 773–784 (1961).

15. Trivedi, P. C., Bartlett, J. J. & Pulinilkunnil, T. Lysosomal Biology and Function: Modern View of Cellular Debris Bin. Cells 9, 1131 (2020).

16. Reif, J. S., Schwartz, A. L. & Fallon, R. J. Low concentrations of primaquine inhibit degradation but not receptor-mediated endocytosis of asialoorosomucoid by HepG2 cells. Exp Cell Res 192, 581–586 (1991).

17. Yoshimori, T., Yamamoto, A., Moriyama, Y., Futai, M. & Tashiro, Y. Bafilomycin A1, a specific inhibitor of vacuolar-type H(+)-ATPase, inhibits acidification and protein degradation in lysosomes of cultured cells. J Biol Chem 266, 17707–17712 (1991).

18. Clague, M. J., Urbé, S., Aniento, F. & Gruenberg, J. Vacuolar ATPase activity is required for endosomal carrier vesicle formation. The Journal of biological chemistry 269, 21–24 (1994).

19. Aniento, F., Gu, F., Parton, R. G. & Gruenberg, J. An endosomal beta COP is involved in the pH-dependent formation of transport vesicles destined for late endosomes. The Journal of cell biology 133, 29–41 (1996).

20. Gu, F., Aniento, F., Parton, R. G. & Gruenberg, J. Functional dissection of COP-I subunits in the biogenesis of multivesicular endosomes. The Journal of cell biology 139, 1183–1195 (1997).

21. Gu, F. & Gruenberg, J. ARF1 regulates pH-dependent COP functions in the early endocytic pathway. The Journal of biological chemistry 275, 8154–8160 (2000).

22. Presley, J. F., Mayor, S., McGraw, T. E., Dunn, K. W. & Maxfield, F. R. Bafilomycin A1 treatment retards transferrin receptor recycling more than bulk membrane recycling. The Journal of biological chemistry 272, 13929–13936 (1997).

23. D’Arrigo, A., Bucci, C., Toh, B. H. & Stenmark, H. Microtubules are involved in bafilomycin A1-induced tubulation and Rab5-dependent vacuolation of early endosomes. European journal of cell biology 72, 95–103 (1997).

24. Mesaki, K., Tanabe, K., Obayashi, M., Oe, N. & Takei, K. Fission of tubular endosomes triggers endosomal acidification and movement. PLOS ONE 6, e19764 (2011).

25. Johnson, L. S., Dunn, K. W., Pytowski, B. & McGraw, T. E. Endosome acidification and receptor trafficking: bafilomycin A1 slows receptor externalization by a mechanism involving the receptor’s internalization motif. MBoC 4, 1251–1266 (1993).

26. Basu, S. K., Goldstein, J. L., Anderson, R. G. & Brown, M. S. Monensin interrupts the recycling of low density lipoprotein receptors in human fibroblasts. Cell 24, 493–502 (1981).

27. Tietze, C., Schlesinger, P. & Stahl, P. Chloroquine and ammonium ion inhibit receptor-mediated endocytosis of mannose-glycoconjugates by macrophages: Apparent inhibition of receptor recycling. Biochemical and Biophysical Research Communications 93, 1–8 (1980).

28. Grant, K. I. et al. Ammonium chloride causes reversible inhibition of low density lipoprotein receptor recycling and accelerates receptor degradation. J Biol Chem 265, 4041–4047 (1990).

29. van Weert, A. W., Dunn, K. W., Geuze, H. J., Maxfield, F. R. & Stoorvogel, W. Transport from late endosomes to lysosomes, but not sorting of integral membrane proteins in endosomes, depends on the vacuolar proton pump. J Cell Biol 130, 821–834 (1995).

30. Bayer, N. et al. Effect of bafilomycin A1 and nocodazole on endocytic transport in HeLa cells: implications for viral uncoating and infection. J Virol 72, 9645–9655 (1998).

31. van Deurs, B., Holm, P. K. & Sandvig, K. Inhibition of the vacuolar H(+)-ATPase with bafilomycin reduces delivery of internalized molecules from mature multivesicular endosomes to lysosomes in HEp-2 cells. Eur J Cell Biol 69, 343–350 (1996).

32. Furuchi, T., Aikawa, K., Arai, H. & Inoue, K. Bafilomycin A1, a specific inhibitor of vacuolar-type H(+)-ATPase, blocks lysosomal cholesterol trafficking in macrophages. Journal of Biological Chemistry 268, 27345–27348 (1993).

33. Burgert, H. G. & Thilo, L. Internalization and recycling of plasma membrane glycoconjugates during pinocytosis in the macrophage cell line, P388D1. Kinetic evidence for compartmentation of internalized membranes. Exp Cell Res 144, 127–142 (1983).

34. Stenmark, H. et al. Inhibition of rab5 GTPase activity stimulates membrane fusion in endocytosis. The EMBO Journal 13, 1287–1296 (1994).

35. Periasamy, A., Mazumder, N., Sun, Y., Christopher, K. G. & Day, R. N. FRET Microscopy: Basics, Issues and Advantages of FLIM-FRET Imaging. in Advanced Time-Correlated Single Photon Counting Applications (ed. Becker, W.) vol. 111 249–276 (Springer International Publishing, 2015).

36. Cantalupo, G., Alifano, P., Roberti, V., Bruni, C. B. & Bucci, C. Rab-interacting lysosomal protein (RILP): the Rab7 effector required for transport to lysosomes. EMBO J 20, 683–693 (2001).

37. Steger, M. et al. Phosphoproteomics reveals that Parkinson’s disease kinase LRRK2 regulates a subset of Rab GTPases. eLife 5, e12813 (2016).

38. Liu, Z. et al. LRRK2 phosphorylates membrane-bound Rabs and is activated by GTP-bound Rab7L1 to promote recruitment to the trans-Golgi network. Human Molecular Genetics 27, 385–395 (2018).

39. Waschbüsch, D. et al. Structural Basis for Rab8a Recruitment of RILPL2 via LRRK2 Phosphorylation of Switch 2. Structure/Folding and Design 28, 406–417.e6 (2020).

40. Waschbüsch, D. & Khan, A. R. Phosphorylation of Rab GTPasesin the regulation of membrane trafficking. Traffic (Copenhagen, Denmark) 21, 712–719 (2020).

41. Sheng, Z. et al. Ser1292 autophosphorylation is an indicator of LRRK2 kinase activity and contributes to the cellular effects of PD mutations. Science translational medicine 4, 164ra161–164ra161 (2012).

42. Gloeckner, C. J. et al. Phosphopeptide Analysis Reveals Two Discrete Clusters of Phosphorylation in the N-Terminus and the Roc Domain of the Parkinson-Disease Associated Protein Kinase LRRK2. Journal of Proteome Research 9, 1738–1745 (2010).

43. Greggio, E. et al. The Parkinson disease-associated leucine-rich repeat kinase 2 (LRRK2) is a dimer that undergoes intramolecular autophosphorylation. The Journal of biological chemistry 283, 16906–16914 (2008).

44. Lagache, T. et al. Mapping molecular assemblies with fluorescence microscopy and object-based spatial statistics. Nature Communications 9, 698–15 (2018).

45. Kuwahara, T. et al. Roles of lysosomotropic agents on LRRK2 activation and Rab10 phosphorylation. Neurobiology of Disease 145, 105081 (2020).

46. Herbst, S. et al. LRRK2 activation controls the repair of damaged endomembranes in macrophages. The EMBO Journal 39, e104494 (2020).

47. Eguchi, T. et al. LRRK2 and its substrate Rab GTPases are sequentially targeted onto stressed lysosomes and maintain their homeostasis. Proceedings of the National Academy of Sciences 115, E9115–E9124 (2018).

48. Bonet-Ponce, L. et al. LRRK2 mediates tubulation and vesicle sorting from lysosomes. Science advances 6, (2020).

49. Kluss, J. H. et al. Detection of endogenous S1292 LRRK2 autophosphorylation in mouse tissue as a readout for kinase activity. NPJ Parkinson’s disease 4, 13 (2018).

50. Fell, M. J. et al. MLi-2, a Potent, Selective, and Centrally Active Compound for Exploring the Therapeutic Potential and Safety of LRRK2 Kinase Inhibition. The Journal of pharmacology and experimental therapeutics 355, 397–409 (2015).

51. Lis, P. et al. Development of phospho-specific Rab protein antibodies to monitor in vivo activity of the LRRK2 Parkinson’s disease kinase. Biochemical Journal 475, 1–22 (2018).

52. Maranda, B. et al. Intra-endosomal pH-sensitive recruitment of the Arf-nucleotide exchange factor ARNO and Arf6 from cytoplasm to proximal tubule endosomes. The Journal of biological chemistry 276, 18540–18550 (2001).

53. Hurtado-Lorenzo, A. et al. V-ATPase interacts with ARNO and Arf6 in early endosomes and regulates the protein degradative pathway. Nature Cell Biology 8, 124–136 (2006).

54. Naufer, A. et al. pH of endophagosomes controls association of their membranes with Vps34 and PtdIns(3)P levels. The Journal of cell biology 55, jcb.201702179 (2017).

55. Sun, L. et al. CED-10/Rac1 Regulates Endocytic Recycling through the RAB-5 GAP TBC-2. PLOS Genetics 8, e1002785 (2012).

56. Jimenez-Orgaz, A. et al. Control of RAB7 activity and localization through the retromer-TBC1D5 complex enables RAB7-dependent mitophagy. EMBO J 37, 235–254 (2018).

57. Haas, A. K., Fuchs, E., Kopajtich, R. & Barr, F. A. A GTPase-activating protein controls Rab5 function in endocytic trafficking. Nature Cell Biology 7, 887–893 (2005).

58. Chotard, L. et al. TBC-2 Regulates RAB-5/RAB-7-mediated Endosomal Trafficking in Caenorhabditis elegans. Molecular Biology of the Cell 21, 2285–2296 (2010).

59. Steger, M. et al. Systematic proteomic analysis of LRRK2-mediated Rab GTPase phosphorylation establishes a connection to ciliogenesis. eLife 6, e31012 (2017).

60. Pfeffer, S. R. LRRK2 and Rab GTPases. Biochemical Society Transactions 46, 1707–1712 (2018).

61. Funayama, M. et al. A new locus for Parkinson’s disease (PARK8) maps to chromosome 12p11.2-q13.1. Ann Neurol 51, 296–301 (2002).

62. Paisán-Ruíz, C. et al. Cloning of the gene containing mutations that cause PARK8-linked Parkinson’s disease. Neuron 44, 595–600 (2004).

63. Zimprich, A. et al. Mutations in LRRK2 cause autosomal-dominant parkinsonism with pleomorphic pathology. Neuron 44, 601–607 (2004).

64. Ross, O. A. et al. Association of LRRK2 exonic variants with susceptibility to Parkinson’s disease: a case-control study. Lancet Neurol 10, 898–908 (2011).

65. Wallings, R., Connor-Robson, N. & Wade-Martins, R. LRRK2 interacts with the vacuolar-type H+-ATPase pump a1 subunit to regulate lysosomal function. Human Molecular Genetics 28, 2696–2710 (2019).

66. Marshansky, V., Rubinstein, J. L. & Grüber, G. Eukaryotic V-ATPase: Novel structural findings and functional insights. Biochimica et biophysica acta 1837, 857–879 (2014).

67. Marshansky, V. & Futai, M. The V-type H+-ATPase in vesicular trafficking: targeting, regulation and function. Current opinion in cell biology 20, 415–426 (2008).

68. Merkulova, M. et al. Structural model of a2-subunit N-terminus and its binding interface for cytohesin-2: Implication for regulation of V-ATPase function. The FASEB Journal (2013) doi:10.1096/fasebj.27.1_supplement.1001.3.

69. Hosokawa, H. et al. The N termini of a-subunit isoforms are involved in signaling between vacuolar H+-ATPase (V-ATPase) and cytohesin-2. The Journal of biological chemistry 288, 5896–5913 (2013).

70. Merkulova, M., Bakulina, A., Thaker, Y. R., Grüber, G. & Marshansky, V. Specific motifs of the V-ATPase a2-subunit isoform interact with catalytic and regulatory domains of ARNO. Biochimica et biophysica acta 1797, 1398–1409 (2010).

71. Gomez, R. C., Wawro, P., Lis, P., Alessi, D. R. & Pfeffer, S. R. Membrane association but not identity is required for LRRK2 activation and phosphorylation of Rab GTPases. J Cell Biol 218, 4157–4170 (2019).

72. Purlyte, E. et al. Rab29 activation of the Parkinson’s disease-associated LRRK2 kinase. The EMBO Journal 37, 1–18 (2018).

73. Healy, D. G. et al. Phenotype, genotype, and worldwide genetic penetrance of LRRK2-associated Parkinson’s disease: a case-control study. Lancet Neurol 7, 583–590 (2008).

74. Domingo, A. & Klein, C. Genetics of Parkinson disease. Handb Clin Neurol 147, 211–227 (2018).

75. Alegre-Abarrategui, J. et al. LRRK2 regulates autophagic activity and localizes to specific membrane microdomains in a novel human genomic reporter cellular model. Human Molecular Genetics 18, 4022–4034 (2009).

76. Manzoni, C. et al. Inhibition of LRRK2 kinase activity stimulates macroautophagy. Biochimica et biophysica acta 1833, 2900–2910 (2013).

77. Rocha, E. M. et al. LRRK2 inhibition prevents endolysosomal deficits seen in human Parkinson’s disease. Neurobiology of Disease 134, 104626 (2020).

78. Jeong, G. R. et al. Dysregulated phosphorylation of Rab GTPases by LRRK2 induces neurodegeneration. Molecular Neurodegeneration 13, 8 (2018).

79. MacLeod, D. A. et al. RAB7L1 interacts with LRRK2 to modify intraneuronal protein sorting and Parkinson’s disease risk. Neuron 77, 425–439 (2013).

80. Orenstein, S. J. et al. Interplay of LRRK2 with chaperone-mediated autophagy. Nat Neurosci 16, 394–406 (2013).

81. Kluss, J. H. et al. Lysosomal positioning regulates Rab10 phosphorylation at LRRK2+ lysosomes. Proceedings of the National Academy of Sciences 119, e2205492119 (2022).

82. Heo, W. D. et al. PI(3,4,5)P3 and PI(4,5)P2 lipids target proteins with polybasic clusters to the plasma membrane. Science 314, 1458–1461 (2006).

83. Roberts, R. L., Barbieri, M. A., Ullrich, J. & Stahl, P. D. Dynamics of rab5 activation in endocytosis and phagocytosis. Journal of Leukocyte Biology 68, 627–632 (2000).

84. Chen, P.-I., Kong, C., Su, X. & Stahl, P. D. Rab5 isoforms differentially regulate the trafficking and degradation of epidermal growth factor receptors. Journal of Biological Chemistry 284, 30328–30338 (2009).

85. D’Costa, V. M. et al. Salmonella Disrupts Host Endocytic Trafficking by SopD2-Mediated Inhibition of Rab7. Cell Reports 12, 1508–1518 (2015).

86. Bucci, C., Thomsen, P., Nicoziani, P., McCarthy, J. & van Deurs, B. Rab7: a key to lysosome biogenesis. Mol Biol Cell 11, 467–480 (2000).

87. Galperin, E. & Sorkin, A. Visualization of Rab5 activity in living cells by FRET microscopy and influence of plasma-membrane-targeted Rab5 on clathrin-dependent endocytosis. Journal of cell science 116, 4799–4810 (2003).

88. Zoncu, R. et al. A phosphoinositide switch controls the maturation and signaling properties of APPL endosomes. Cell 136, 1110–1121 (2009).

89. Phair, R. D., Gorski, S. A. & Misteli, T. Measurement of dynamic protein binding to chromatin in vivo, using photobleaching microscopy. Methods in enzymology 375, 393–414 (2004).

90. Kang, M., Day, C. A., Kenworthy, A. K. & DiBenedetto, E. Simplified equation to extract diffusion coefficients from confocal FRAP data. Traffic (Copenhagen, Denmark) 13, 1589– 1600 (2012).

